# MAVS mediates a protective immune response in the brain to Rift Valley fever virus

**DOI:** 10.1101/2021.12.22.473954

**Authors:** Nicholas R. Hum, Feliza A. Bourguet, Aimy Sebastian, Doris Lam, Ashlee M. Phillips, Kristina R. Sanchez, Amy Rasley, Gabriela G. Loots, Dina R. Weilhammer

## Abstract

Rift Valley fever virus (RVFV) is a highly pathogenic mosquito-borne virus capable of causing hepatitis, encephalitis, blindness, hemorrhagic syndrome, and death in humans and livestock. Upon aerosol infection with RVFV, the brain is a major site of viral replication and tissue damage, yet pathogenesis in this organ has been understudied. Here, we investigated the immune response in the brain of RVFV infected mice. In response to infection, microglia initiate robust transcriptional upregulation of antiviral immune genes, as well as increased levels of activation markers and cytokine secretion that is dependent on mitochondrial antiviral-signaling protein (MAVS) and independent of toll-like receptors 3 and 7. *In vivo, Mavs*^*-/-*^ mice displayed enhanced susceptibility to RVFV as determined by increased brain viral burden and higher mortality. Single-cell RNA sequence analysis identified microglia-specific defects in type I interferon and interferon responsive gene expression in *Mavs*^*-/-*^ mice, as well as dysregulated lymphocyte infiltration. The results of this study provide a crucial step towards understanding the precise molecular mechanisms by which RVFV infection is controlled in the brain and will help inform the development of vaccines and antiviral therapies that are effective in preventing encephalitis.

**Author Summary:** Rift Valley fever virus causes severe disease in humans and livestock and in some cases can be fatal. There is concern about the use of Rift Valley fever virus as a bioweapon since it can be transmitted through the air, and there are no vaccines or antiviral treatments. Airborne transmission of the virus causes severe inflammation of the brain, yet little is known about the immune response against the virus in this organ. Here, we investigated the immune response in the brain to Rift Valley fever virus following intranasal infection. We determined that microglia, the resident immune cells of the brain, initiate a robust response to Rift Valley fever virus infection and identified a key immune pathway that is critical for the ability of microglia to respond to infection. When this immune pathway is rendered non-functional, mice have a dysregulated response to infection in the brain. This study provides insight into how the immune response can control Rift Valley fever virus infection of the brain.

## Introduction

Rift Valley fever virus (RVFV) (genus *Phlebovirus/*family *Bunyaviridae*), is a highly pathogenic mosquito-borne virus that can cause lethal disease in both humans and livestock, including acute-onset hepatitis, delayed-onset encephalitis, blindness, or hemorrhagic fever [1]. RVFV is endemic to Africa however concern exists about its potential spread across the world, similar to Zika or West Nile virus (WNV) [1]. Outside of Africa, competent vectors for RVFV include over 30 species of mosquitoes that are present throughout North and South America [2]. RVFV is also classified as a Category A Biodefense pathogen by the National Institute of Allergy and Infectious Diseases (NIAID) due to the potential for intentional spread by aerosol and the lack of licensed vaccines or antiviral therapeutics. Furthermore, RVFV is classified by the Department of Health and Human Services (HHS) and United States Department of Agriculture (USDA) as an overlap select agent due to the susceptibility of numerous livestock species to this disease [3].

In the event of an intentional release, the most likely route of exposure to RVFV would be through the respiratory system via aerosolized release of the virus. Studies conducted using rodent models indicate a more severe infection following respiratory versus subcutaneous exposure, with higher incidence of lethality, neuropathology, and increased viral titers in the brain [4-7]. Analysis of human infections also suggests that aerosol exposure to RVFV (i.e., laboratory acquired infections and infections acquired via handling of infected livestock) leads to a higher incidence of severe disease with encephalitis and long-lasting neurologic complications [8, 9]. The fatality rate amongst patients with encephalitic manifestations of disease is ∼50% [10], which is much higher than the overall fatality rate, estimated to be between 0.5-2% [11]. Thus, a thorough understanding of RVFV pathogenesis in the brain is required for preparedness to combat the virus’ worst outcomes, including intentional release of the virus.

The term “immune privileged site” was once applied to the brain and interpreted to mean that few immune defenses were functional in this organ [12]. However, it is now widely accepted that the brain is highly immunologically active [13]. An emerging body of evidence indicates that immune responses within the brain are critical for control of an array of neuroinvasive viruses [14, 15]. In the brain, neurons, and glial cells (e.g., astrocytes, and microglia), express many of the same pattern recognition receptors (PRRs) expressed by cells in the periphery and initiate type I interferon (IFN) expression as well as other innate responses upon viral infection [15, 16]. Such early responses in the brain are critical for direct control of viral replication, as well as for recruitment of adaptive immune cells that participate in viral clearance [17, 18]. Microglia, the resident immune cells of the brain, play a key role in bridging innate and adaptive immune responses in the brain [19-21], and depletion of microglia increases susceptibility to multiple viral infections [18, 22, 23]. Although potent immune responses are required for viral control in the brain, limiting inflammation presents a unique immunoregulatory challenge as excessive inflammation can be especially deleterious and promote neurodegenerative diseases such as Parkinson’s [24] and Alzheimer’s [25] disease. Thus, investigation of immune responses in the brain presents an opportunity to understand the interaction of processes that hone the correct response to control viral replication without inducing excessive damaging inflammation.

To date, there has been limited investigation of the immune response to RVFV in the brain. Previous work has indicated that a strong adaptive response involving both CD4 and CD8 T cells as well as a robust antibody response is required for the prevention of encephalitic disease [7, 26, 27] yet there remains a lack of understanding of the response of resident and infiltrating immune cells in the brain. Furthermore, the PRRs that microglia use to respond to RVFV infection have not yet been identified. Previous work has demonstrated the critical role of RIG-I-like receptor (RLR) signaling *via* mitochondrial antiviral signaling (MAVS) in the type I IFN response of macrophages and dendritic cells (DCs) to RVFV infection, as well as a protective role for MAVS following *in vivo* challenge, with little to no contribution from the RNA-sensing toll-like receptors (TLRs) [6]. However, differential roles for the RNA-sensing PRRs during viral infections of the CNS have been identified, most notably for WNV, where TLR3, TLR7, and RLR receptors RIG-I and MDA-5 have been shown to coordinate and propagate a protective response [28-30]. Identifying the relative contributions of innate signaling pathways to the induction of type I IFNs and subsequent control of viral infection has important consequences in terms of understanding human susceptibility to RVFV infection, as polymorphisms in TLR3, TLR7, their respective downstream signaling adaptors TRIF and MyD88, as well as RIG-I and MAVS have all been associated with severe disease/neuropathology in humans [31].

Here, we investigated the immune response in the murine brain to RVFV intranasal infection. We demonstrate that microglia mount a robust response that is dependent on MAVS and independent of TLR3 and TLR7. MAVS is critical for the expression of immunoregulatory genes, secretion of cytokines, and upregulation of surface markers of activation. *Mavs*^*-/-*^ mice are more susceptible to infection, with higher viral titers in the brain following intranasal challenge. RNA sequence (RNA-seq) analysis of whole brain tissue revealed robust immune gene expression with greater induction of inflammatory genes in the brains of *Mavs*^*-/-*^ versus wild type (WT) mice. Single cell RNA-sequence (scRNA-seq) analysis revealed defects in specific antiviral genes and signaling pathways within microglia in the brains of RVFV infected *Mavs*^*-/-*^ mice. The lack of MAVS resulted in a shift towards a more inflammatory phenotype, with a decrease in antiviral signaling pathways and an increase in proinflammatory pathways within *Mavs*^*-/-*^ microglia. Differences in immune infiltration into the brain were also observed between WT and *Mavs*^*-/-*^ mice. These results are an important step towards understanding the cell types and molecular pathways responsible for controlling RVFV infection in the brain and towards future developments of antiviral treatments.

## Results

### RVFV infection induces a robust response in microglia that is dependent upon Mavs and independent of Tlr3 and Tlr7

To determine if microglia respond to RVFV infection directly *via* cell intrinsic mechanisms, we first confirmed infection of microglia cell lines EOC 13.31 and SIM A9, as well as primary microglia derived from the brains of neonatal mice *via* flow cytometry using a RVFV-specific antibody (Figure 1A and B). Vero cells were also included as a positive control for infection. EOC 13.31 cells had the lowest infectivity rate ranging from 30-40% positive cells, while primary microglia displayed infectivity rates (50-70%) similar to SIM A9 cells (60-70%) but slightly less effective than the Vero positive control (90%) (Figure 1B). Both the fully virulent (ZH501) and attenuated (MP-12) strains of RVFV yielded similar levels of infected primary microglia (Figure 1C). Next, we probed the response of primary microglia to both the ZH501 and MP-12 strains by quantifying changes in expression of genes involved in antiviral immune responses using real-time reverse transcription (RT^2^) PCR array (Figure 1D and Table S1). There was a significant response to both viruses with similar patterns of gene expression changes, although the overall magnitude of the response was greater in microglia infected with the attenuated versus the fully virulent strain, consistent with prior reports [32]. The response to MP-12 was robust, with very high levels of *Ifnb1* expression (greater than 7,000-fold upregulation) and greater than 10-fold upregulation of over 20 genes, including *Ifna2, Ifih1, Isg15, Cxcl10, Il6*, and *Il12b*, among others.

**Figure 1.**
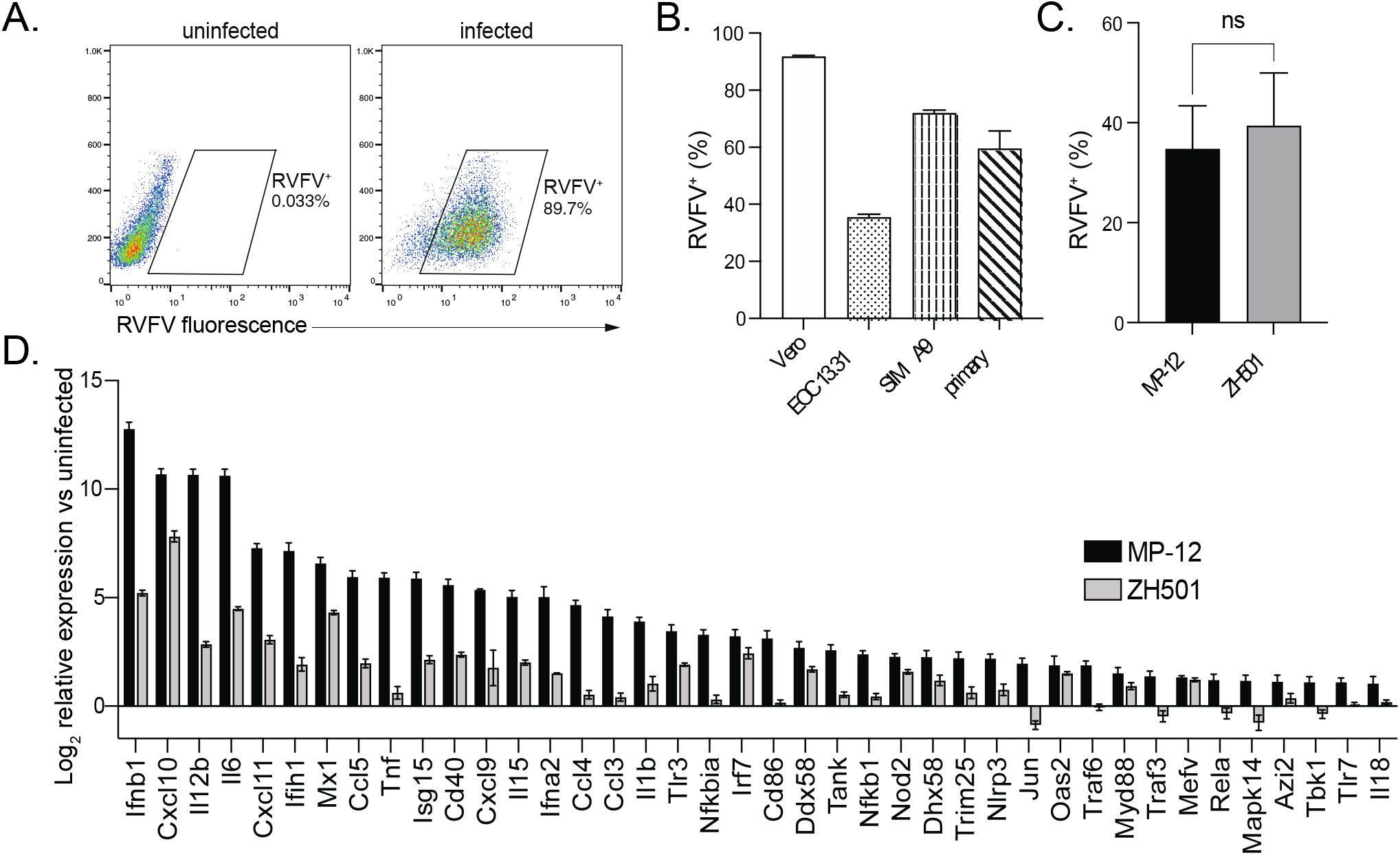
RVFV infects microglia. Microglia cell lines EOC 13.31 and SIM A9 and primary microglia derived from the brains of neonatal mice, along with Vero positive control cells, were infected with RVFV MP-12. The percentage of cells positive for RVFV was assessed by flow cytometry. Representative plots of Vero cell infection analysis (A). RVFV infection rate for all cell types (B). Primary microglia derived from wildtype (WT) mice were infected with RVFV MP-12 or ZH501 and the percentage of cells positive for RVFV was assessed by flow cytometry (C). The relative expression of 40 antiviral response genes with the highest fold change in infected versus uninfected cells (D). Data in (B-C) are shown as the mean +/- SD. Data in (D) are shown as the mean log_2_ fold change in infected versus uninfected cells +/- SEM.

Next, microglia derived from WT, *Tlr3*^*-/-*^, *Tlr7*^*-/-*^, or *Mavs*^*-/-*^ mice were infected with MP-12, and a similar infectivity rate was confirmed amongst all groups, therefore was independent of genotype (Figure S1). Microglia derived from TLR3 or TLR7 deficient animals activated immune genes in response to RVFV at levels similar to WT microglia (Figure 2A and B and Table S1). In contrast, microglia derived from *Mavs*^*-/-*^ mice displayed an abrogated response to infection, with minimal changes in gene expression in most genes in the array and a greater than 4,000-fold reduction in *Ifnb1* expression versus WT infected microglia (Figure 2C and Table S1).

**Figure 2.**
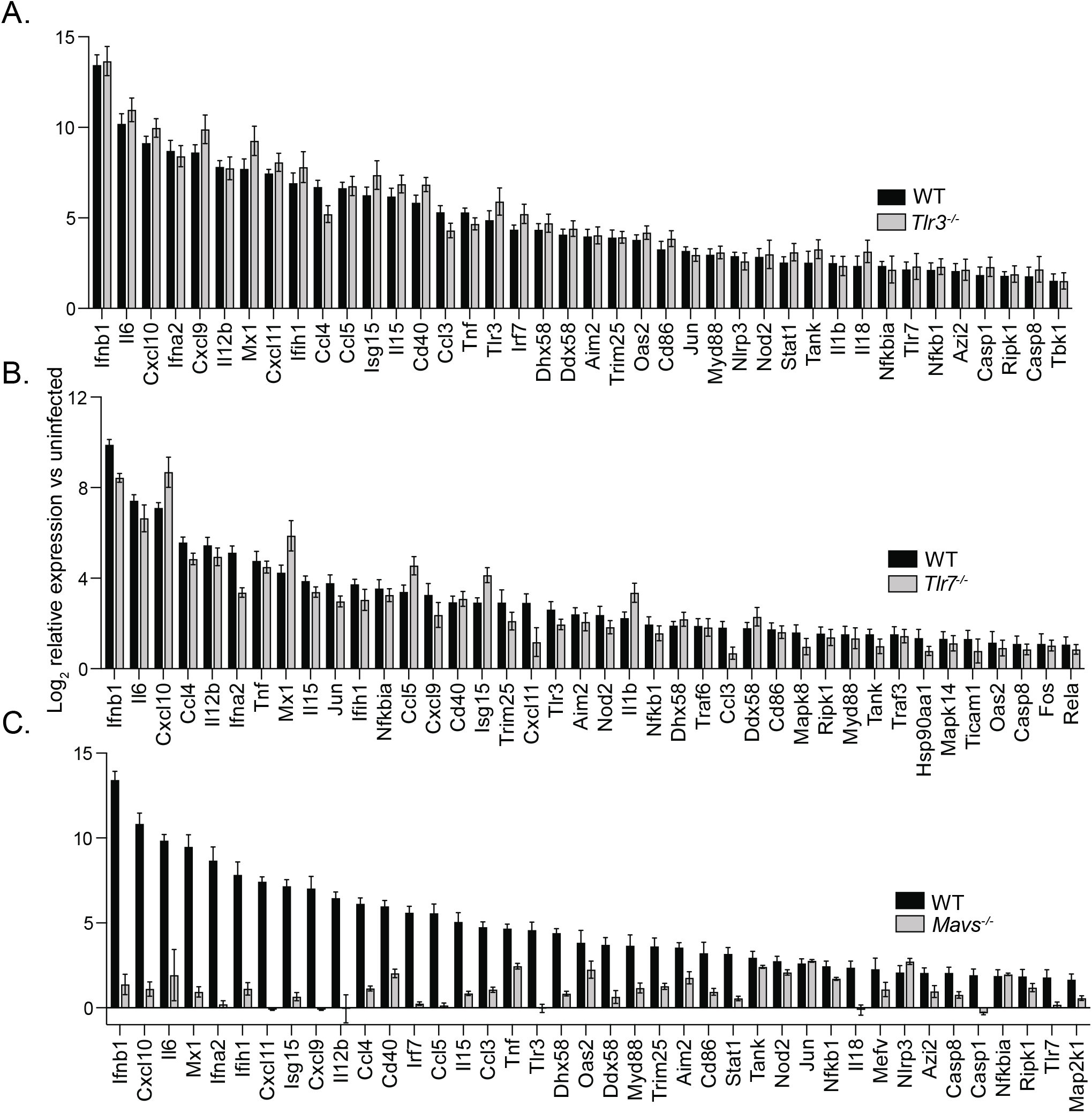
Microglial response to RVFV infection is dependent upon MAVS and independent of TLR7 and TLR3. Primary microglia derived from WT or *Tlr3*^*-/-*^ (A), *Tlr7*^*-/-*^ (B) or *Mavs*^*-/-*^ (C) were infected with RVFV MP-12. The relative expression of 40 antiviral response genes with the highest fold change in WT cells are shown as the mean log_2_ fold change in infected versus uninfected cells, +/- SEM.

We further characterized the role of MAVS in the response of microglia to RVFV infection *in vitro* by assessing expression of microglial activation markers as well as cytokine secretion (Figure 3). Microglia upregulated surface expression of CD86, CD80, and I-A/I-E in a MAVS-dependent manner (Figure 3A). Using RVFV antibody staining, cells within infected cultures could be identified as infected (RVFV^+^) or uninfected (RVFV^-^). Expression was upregulated not only on infected microglia (RVFV^+^), but also on uninfected cells within the culture (RVFV^-^), suggesting secreted cytokines can influence the activation state of cells in trans. Within WT microglia, expression of each marker was highest on infected cells, with an intermediate level of expression seen on cells activated in trans. On *Mavs*^*-/-*^ microglia, apart from a low level of CD80 on infected cells, no significant upregulation of activation markers was observed. In contrast, upregulation of activation markers was unaltered from WT on *Tlr3*^*-/-*^ and *Tlr7*^*-/-*^ microglia (Figure S2).

**Figure 3.**
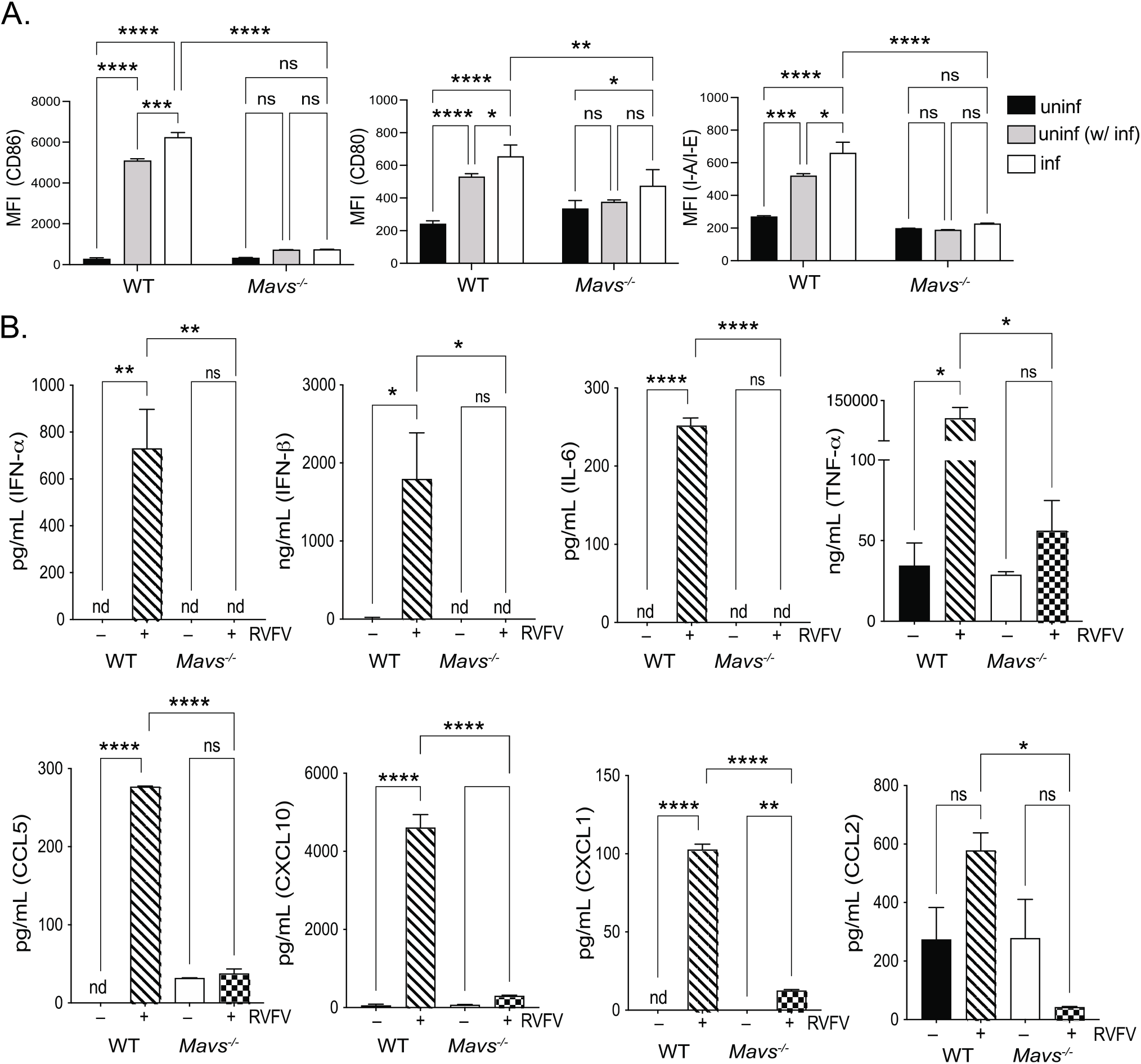
Microglia respond to RVFV infection in vitro by cytokine secretion and upregulation of activation markers. Microglia derived from WT or *Mavs*^*-/-*^ mice were infected with RVFV MP-12 and at 18-24 h post-infection, cells and cellular supernatants were harvested for flow cytometry and multiplex cytokine analysis, respectively. The expression levels of the indicated activation markers were assessed on uninfected cells (black bars), uninfected cells in culture with infected cells (uninf (w/ inf), gray bars), and infected cells (inf, white bars) and shown as the mean fluorescence intensity of the indicated activation markers (A). Cytokine levels in cellular supernatants (B). Data are shown as the mean +/- SD, nd = not detected and was denoted as zero *p <0.05, **p<0.01, ***p<0.001, ****p<0.0001

Cytokine secretion by infected microglia was also dependent on MAVS (Figure 3B). High levels of type I IFNs were detected in supernatants from WT infected cells, as well the inflammatory cytokines IL-6 and TNF-α, and chemokines CCL5, CXCL10, CXCL11, and CCL2. Cytokine levels in supernatants from MAVS-deficient cells were either not detectable, or not significantly different from uninfected supernatants. Taken together, these data demonstrate that microglia have a robust response to RVFV infection that is mediated primarily through the RLR signaling adaptor MAVS.

### Mavs^-/-^ animals have a dysregulated immune response with increased susceptibility to infection

We then investigated the role of MAVS in the immune response in the brain during *in vivo* challenge with RVFV (Figure 4). Intranasal challenge of WT and *Mavs*^*-/-*^ cohorts of mice (n=20) with the ZH501 strain of RVFV confirmed enhanced susceptibility to infection of *Mavs*^*-/-*^ mice, consistent with prior reports [6]. *Mavs*^*-/-*^ mice succumbed to challenge significantly faster, with 100% of *Mavs*^*-/-*^ mice deceased by 6 days post infection, whereas 20% of WT animals were still alive 10 days post infection (Figure 4A). To capture early innate antiviral responses, we conducted a longitudinal study over 4 days, where infected brains of WT mice were examined *ex vivo* to determine when live virus (MP-12) could be cultured from brain tissue following intranasal challenge (Figure S3). Day 7 post-infection was identified as the earliest timepoint live virus could be detected in infected WT brains. Next, we compared viral levels at day 7 post infection with MP-12 between WT and *Mavs*^*-/-*^ mice and were able to detect infection in 100% of *Mavs*^*-/-*^ (n=11), but only in 42% of WT (n=12) infected brains. Within brains with detectable virus, titers of live virus as well as the levels of RVFV genomic RNA were significantly higher in *Mavs*^*-/-*^ than in WT mice (Figure 4B).

**Figure 4.**
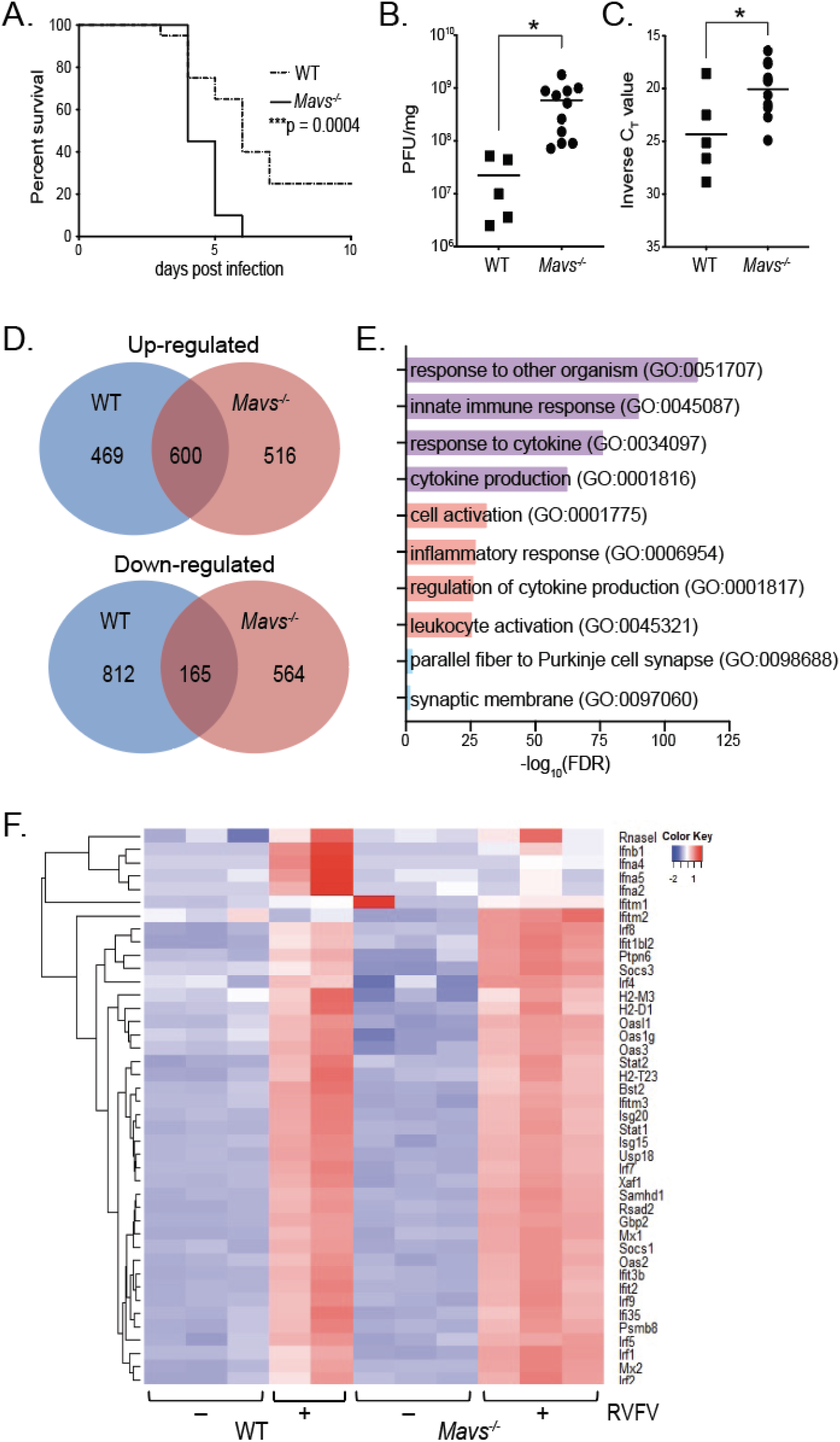
*Mavs*^*-/-*^ mice are more susceptible to intranasal RVFV infection and display increased viral burden and immune gene expression in the brain. WT or *Mavs*^*-/-*^ mice were infected intranasally with 1000 PFU RVFV ZH501 (A), or 5×10^5^ PFU RVFV MP-12 (B-E). Morbidity and mortality were assessed daily, and percent survival is depicted in (A). On day 7 post infection (B-E), brains were processed for viral quantitation and RNA extraction. Viral quantitation (B). Genes differentially expressed between uninfected and infected brains (C). Functional enrichment of differentially expressed genes (D). Ontologies that were enriched in both WT and *Mavs*^*-/-*^ infected brains are highlighted in purple. Pathways enriched only in *Mavs*^*-/-*^ or WT infected brains are highlighted in red and blue, respectively. Heatmap of IFN α/β signaling genes (E). *p<0.05, ***p<0.001

To assess the antiviral response, we utilized RNA-seq to profile the transcriptomic alterations induced by RVFV infection in the brains of WT and *Mavs*^*-/-*^ mice. Infection resulted in significant upregulation of 1,069 genes in the brains of WT mice and 1,116 genes in the brains of *Mavs*^*-/-*^ mice where 600 upregulated genes were in common to both WT and *Mavs*^*-/-*^ (Figure 4C and Table S2). Downregulation of genes in response to infection displayed less overlap between WT and *Mavs*^*-/-*^, with 977 downregulated in WT, 729 downregulated in *Mavs*^*-/-*^, and only 165 downregulated genes common to both genotypes (Figure 4C). Functional enrichment of differentially expressed genes revealed strong correlation of pathways related to innate immune and defense responses in both WT and *Mavs*^*-/-*^ brains, including response to other organism (GO:0051707), innate immune response (GO:0045087), response to cytokine (GO:0034097), and cytokine production (GO:0001816) (Figure 4D).

To identify deficiencies in the antiviral response within *Mavs*^*-/-*^ brains that would indicate the effector functions downstream of MAVS signaling that control viral replication, we focused on pathways that were enriched in WT and unaffected in *Mavs*^*-/-*^ infected brains. Interestingly, only two pathways were enriched in WT and not *Mavs*^*-/-*^ yet neither are involved in immune responses nor indicate any functional advantage WT animals have at controlling viral replication (Figure 4D). *Mavs*^*-/-*^ brains had more exclusively enriched pathways, including regulation of cytokine production (GO:0001817) and inflammatory response (GO:0006954), indicating that the overall levels of inflammation were higher in *Mavs*^*-/-*^ brains than in WT. This is consistent with previous reports that indicated a more robust serum cytokine response in *Mavs*^*-/-*^ than WT mice infected with RVFV [6].

IFN α/β gene expression levels were elevated in WT infected brains, whereas induction was not seen in *Mavs*^*-/-*^ brains (Figure 4E). However, global upregulation of type I interferon signaling was not observed in the whole brain between WT and *Mavs*^*-/-*^. Thus, while a deficiency in type I interferons is noted in *Mavs*^*-/-*^ brains, examination of downstream signaling does not reveal potential deficiencies in antiviral response genes. In total, the data demonstrate that RVFV infection results in robust activation of immune and inflammatory genes in both WT and *Mavs*^*-/-*^ brains with very similar patterns of gene upregulation in both.

### Single cell RNA sequencing reveals changes in immune infiltration and signaling defects in microglia following RVFV infection of Mavs^-/-^ mice

Next, we performed scRNA-seq to increase the resolution of transcriptional analysis and assess cellular variations resulting from the absence of MAVS during RVFV infection of the brain. Single cell suspensions were generated from whole brain tissue of WT and *Mavs*^*-/-*^ mice, with (WT+, *Mavs*^*-/-*^+) and without (WT-, *Mavs*^*-/-*^-) RVFV infection. The following cell numbers were sequenced from each condition: WT-: 1,989, *Mavs*^*-/-*^-: 1,749, WT+: 1627, *Mavs*^*-/-*^-: 1841, for a total of 7,206 cells. Unsupervised clustering of the data resulted in 15 cell type clusters (Figure 5A). By cross-referencing genes differentially expressed in each cluster to previously published cell-type specific markers [33-36], we assigned each cluster to its putative cell-type identity (Figure 5C). We identified expected cell clusters such as neurons, astrocytes, oligodendrocytes, and endothelial cells. Three clusters of immune cells were identified, corresponding to microglia/myeloid cells, T/natural killer (NK) cells, and neutrophils. We observed a large shift in the relative frequency of immune cells upon infection, indicating massive immune infiltration in the brains of infected animals (Figure 5B and D). Sequenced cells from uninfected brains were comprised of 70-80% non-immune cells (clusters 1, 2, 4-9, and 11), with immune cells (clusters 0, 3 and 10) comprising less than 30%. In contrast, sequenced cells from infected brains were comprised of 30-40% non-immune and more than 60% immune cells (Figure 5B and D). Interestingly, the pattern of immune infiltration differed between WT and *Mavs*^*-/-*^ infected brains. WT animals exhibited infiltration of mostly myeloid lineage cells, whereas *Mavs*^*-/-*^ animals exhibited infiltration of lymphocyte populations in addition to myeloid lineage cells (Figure 5D). Immune infiltration was confirmed using flow cytometry (Figure 5 E-G) and was consistent with scRNA-seq ratios. Lymphocytes and non-glial cells of myeloid lineage comprised a very small fraction of immune cells within uninfected brains, ranging from 0.5-4% and 1-6% of brain immune cells, respectively (Figure 5F and G). In contrast, the percentage of non-glial myeloid cells increased to 10-25% of brain immune cells in both WT and *Mavs*^*-/-*^ infected mice (Figure 5F). Lymphocytes also increased in the brains of both WT and *Mavs*^*-/-*^ infected mice, however significantly more lymphocytes were observed in the brains of *Mavs*^*-/-*^ mice (Figure 5G), with lymphocytes comprising 5-15% of brain immune cells in WT mice and 15-30% of brain immune cells in *Mavs*^*-/-*^ mice.

**Figure 5.**
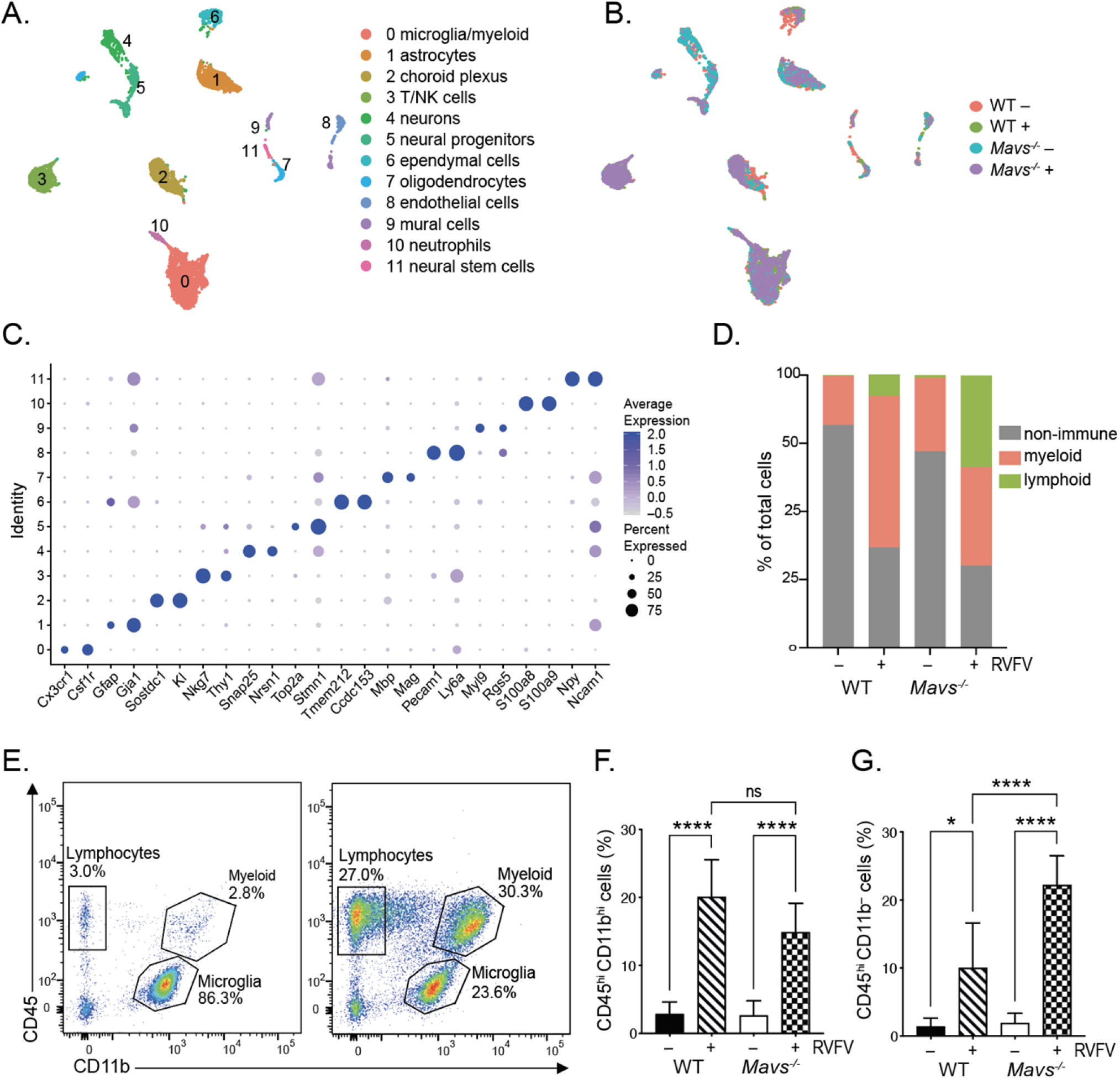
Single cell RNA sequencing reveals shifts in cell populations and immune infiltration in the brain following RVFV infection. UMAP plots depicting cell clusters derived from WT and *Mavs*^*-/-*^ brains, +/- RVFV infection. Cell types (A) or conditions (B) are color coded. Gene markers for specific cell types (C). The distribution of cell types within brains of each condition (D). Flow cytometry: Representative plots of CD11b vs CD45 and the gates defining microglia, lymphocytes, and myeloid cells are shown for uninfected (left panel) and infected brains (right panel) (E). The percent myeloid cells (CD11b^hi^ CD45^hi^), and lymphocytes (CD11b^-^ CD45^hi^) (F) and (G), respectively. Data in (F) and (G) are shown as the mean +/- SD. *p<0.05, ***p<0.001, ****p<0.0001

Clustering of microglia/myeloid cells (cluster 0, Figure 5A) identified 6 subclusters of myeloid cells (Figure 6A). Enumeration of cells according to condition suggested that most immune cells present in uninfected brains were microglia, whereas infected brains contained cells of multiple infiltrating lineages including monocytes, antigen presenting cells (APCs), and granulocytic cells (Figure 6B and D). The distribution of myeloid lineage cells in uninfected brains confirmed the flow cytometry data in Figure 5E and is consistent with previously published reports [37] which indicate that microglia are the predominant immune cell type in the brain under non-pathological conditions. The relative frequency of macrophages (cluster 4) was consistent across conditions and likely corresponds to resident non-parenchymal macrophages [38], rather than an infiltrated population. Infected brains of both genotypes demonstrated similar patterns of monocyte, APC, and granulocytic cell infiltration (clusters 2, 3, and 5, respectively). Overall, the distribution of myeloid cells was similar between infected brains of both genotypes.

**Figure 6.**
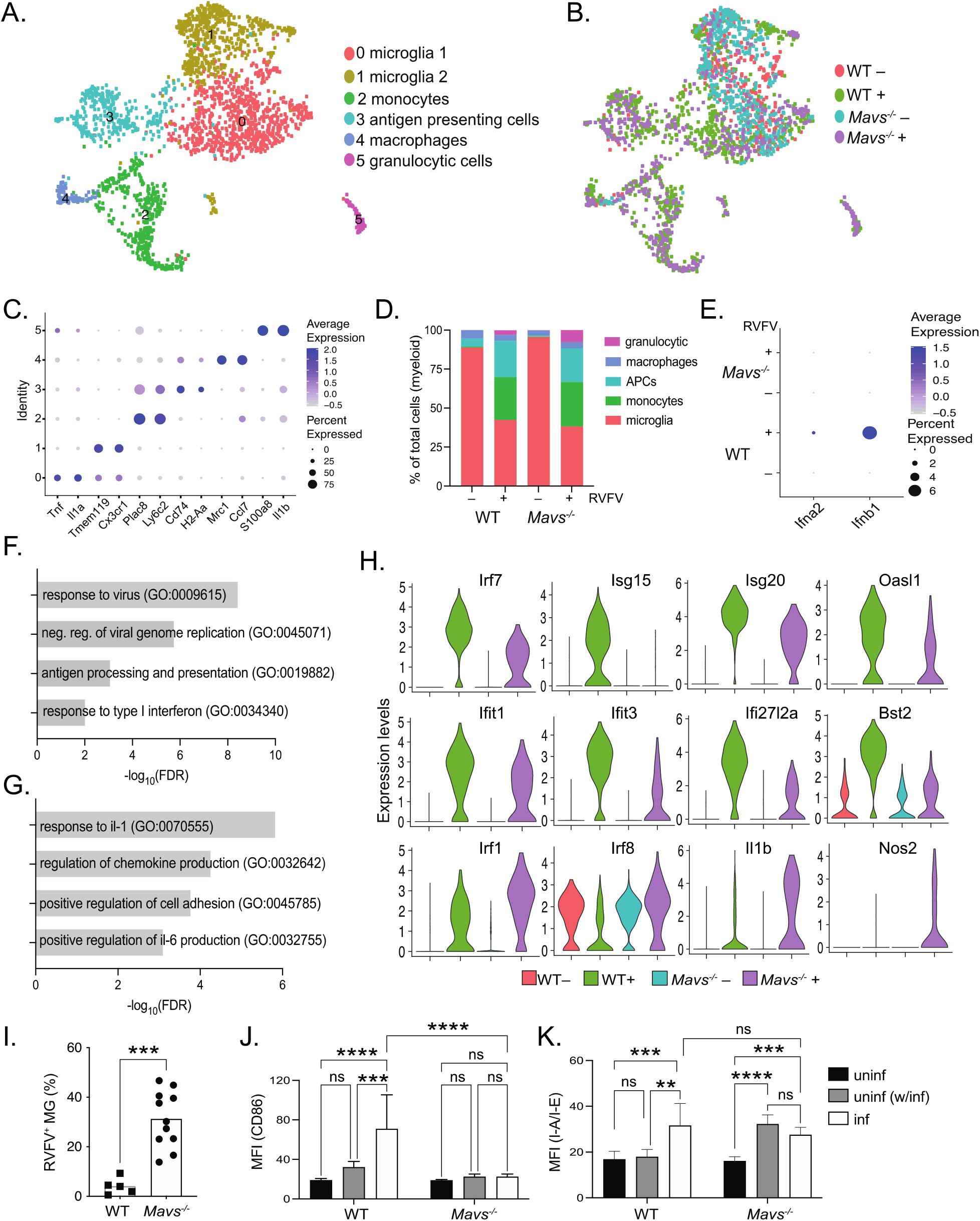
Microglia from *Mavs*^*-/-*^ brains have defective response to RVFV infection. UMAP plots depicting cell clusters of myeloid lineage cells derived from WT and *Mavs*^*-/-*^ brains, +/- RVFV infection. Cell types (A) or conditions (B) are color coded. Gene markers for specific cell types (C). The distribution of myeloid lineage cell types within brains of each condition (D). Expression of type I IFNs within microglia (clusters 0 and 1) (E). Gene ontology (GO) enrichment analysis showing enriched in WT vs *Mavs*^*-/-*^, or *Mavs*^*-/-*^ vs WT microglia (clusters 0 and 1) from infected brains are shown in (F) and (G), respectively. Violin plots depicting the relative expression of selected genes within microglia (clusters 0 and 1) are shown in (H). Flow cytometry: microglia were defined as CD11b^int^ CD45^int^ as described in Figure 5, and the percentage of RVFV^+^ microglia are shown in (I). Mean fluorescence intensities (MFI) of CD86 and I-A/I-E expression on microglia (J) and (K), respectively. Data in (I-K) are shown as the mean +/- SD. **p<0.01, ***p<0.001, ****p<0.0001

Next, we examined transcriptional differences in the antiviral responses of microglia derived from WT and *Mavs*^*-/-*^ infected brains. Differences in type I IFN (*Ifnb1* and *Ifna2*) expression were noted in a subset of microglia in WT infected brains whereas levels were undetectable in microglia from *Mavs*^*-/-*^ infected and uninfected brains of both genotypes (Figure 6E). Gene ontological analysis revealed specific enrichment of pathways involved in antiviral responses including response to virus (GO:009615), negative regulation of viral genome replication (GO:0045071), antigen processing and presentation (GO:0019882), and response to type I interferon (GO:0034340) within WT infected microglia (Figure 6F). In contrast, inflammatory pathways such as response to interleukin-1 (GO:007055) and positive regulation of interleukin-6 production (GO:0032755), as well as pathways involved in cell migration such as regulation of chemokine production (GO:0032642) and positive regulation of cell adhesion (GO:0045785) were enriched in *Mavs*^*-/-*^ microglia (Figure 6G). Specific IFN-stimulated genes (ISGs) were expressed at higher levels within WT microglia, including *Irf7, Isg20, Isg15, Oasl1, Ifit1, Ifit3, Ifi27l2a*, and *Bst2* (Figure 6H). Genes that have been previously associated with a pro-inflammatory state [39-41] were expressed at higher levels within *Mavs*^*-/-*^ microglia, including *Irf1, Irf8, Il1b*, and *Nos2* (Figure 6H).

Flow cytometric analysis revealed further defects in *Mavs*^*-/-*^ microglia. In line with increased viral titers in the brains of *Mavs*^*-/-*^ animals (Figure 4B), microglia from *Mavs*^*-/-*^ animals displayed dramatically higher levels of infection as evidenced by anti-RVFV antibody staining, ranging from 0.5-9% in WT brains to 15-50% in *Mavs*^*-/-*^ brains (Figure 6I). We also evaluated the surface expression levels of activation markers; expression of CD86 mirrored *in vitro* results, with WT microglia that are RVFV^+^ expressing the highest levels of CD86, and microglia from infected brains that are RVFV^-^ expressing an intermediate level of CD86, although not significantly different from microglia from uninfected brains (Figure 6J). There was no significant upregulation of CD86 on microglia from *Mavs*^*-/-*^ brains. Expression of I-A/I-E was also elevated on RVFV^+^ microglia, although not on RVFV^-^ microglia, from WT infected brains (Figure 6K). In contrast to *in vitro* results, upregulation of I-A/I-E was detected on microglia derived from *Mavs*^*-/-*^ brains (Figure 6K). No significant upregulation of CD80 was detected (Figure S4).

### Lymphocyte infiltration and signaling defects in the brains of Mavs^-/-^ mice

The majority of lymphocytes identified within WT infected brains were NK cells, whereas those in *Mavs*^*-/-*^ infected brains included a large number of T cells in addition to NK cells (Figure 7A-D). Lymphocytes comprised greater than 30% of the total sequenced cells in the *Mavs*^*-/-*^ infected condition (Figures 5D and 7D). Upon examining genes that were differentially expressed between WT and *Mavs*^*-/-*^ lymphocytes, we observed defects in ISG expression (*Irf7, Ifi44, Xcl1*), as well as genes involved in T and NK cell-mediated killing (*Prf1, Gzmb, Ifng*) [42] within lymphocytes from *Mavs*^*-/-*^ infected animals (Figure 7E), suggesting that *Mavs*^*-/-*^ cells are less efficient in clearing the virus than WT cells [43-45]. Consistent with this, we observed reduced expression of *Il15, Il18*, and *Il27*, cytokines that have a stimulatory effect on T and NK cells [46-49], in microglia from infected *Mavs*^*-/-*^ mice (Figure S5). Lower levels of IFN-γ protein were also detected in the brains of *Mavs*^*-/-*^ versus WT infected mice (Figure 7G). Taken together, these results suggest that a defective response of *Mavs*^*-/-*^ microglia may lead to the reduced capacity of T and NK cells to control RVFV infection.

**Figure 7.**
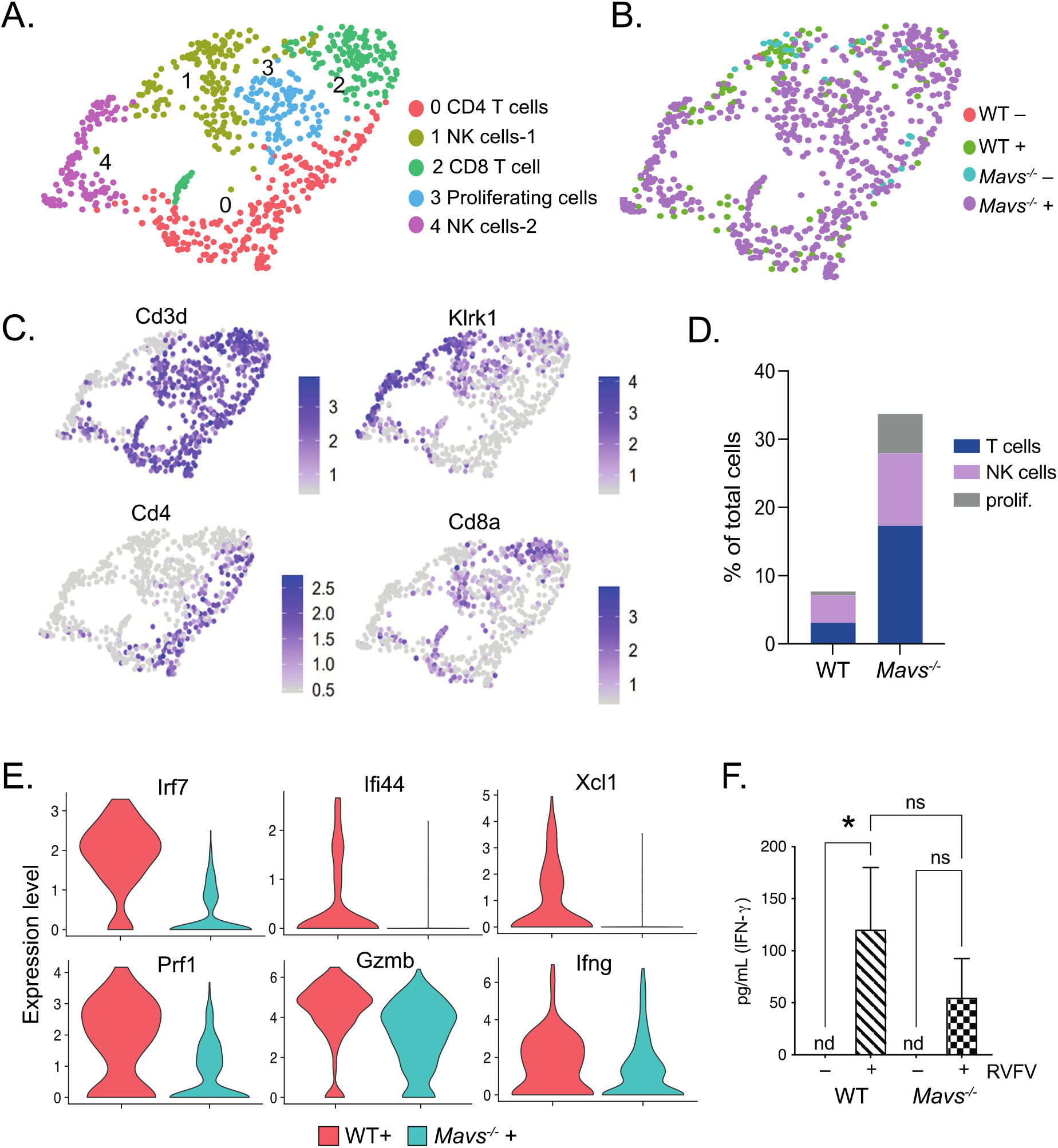
Dysregulated pattern of lymphocyte infiltration and gene expression in brains of *Mavs*^*-/-*^ mice. UMAP plots depicting cell clusters of lymphoid lineage cells derived from WT and *Mavs*^*-/-*^ brains, +/- RVFV infection. Cell types (A) or conditions (B) are color coded. Feature plots showing expression of cell type specific markers are shown in (C). The distribution of lymphoid lineage cell types within brains of infected mice (D). Violin plots depicting the relative expression of selected genes within all lymphocytes are shown in (E). Quantitation of IFN-γ is depicted in (F). *p<0.05

## Discussion

Despite the major role of pathogenesis in the brain for the most severe outcomes of RVFV infection, there has been little investigation of local antiviral immune responses within this organ. Here, we investigated the innate immune response in the brain to RVFV in a mouse model of intranasal infection. We demonstrated that microglia mount a robust response to RVFV that is dependent on MAVS and independent of TLR3 and TLR7. To probe viral pathogenesis in the brain in the presence and absence of a functional innate immune response, we profiled the brains of WT and *Mavs*^*-/-*^ animals following intranasal infection with RVFV. *Mavs*^*-/-*^ animals succumbed to infection more rapidly and with significantly higher viral titers in the brain. Viral presence in the brain corresponded with massive immune infiltration in both WT and *Mavs*^*-/-*^ mice, consisting mostly of myeloid lineage cells as well as some lymphocytes, with significantly more T and NK cells in the brains of *Mavs*^*-/-*^ animals. Robust immune gene expression was observed in the brains of both WT and *Mavs*^*-/-*^ infected animals, with greater inflammatory gene expression within *Mavs*^*-/-*^ brains. Deficiencies in type I IFN expression were noted within whole brain tissue as well as specifically within microglia in *Mavs*^*-/-*^ infected animals, and furthermore, microglia from *Mavs*^*-/-*^ animals displayed deficiencies in downstream antiviral signaling pathways and specific ISG expression. T and NK cells from *Mavs*^*-/-*^ animals were also deficient in ISG expression, as well as genes related to killing functions. Furthermore, decreased *Ifng* expression resulted in lower levels of IFN-γ protein in the brain.

A summary of previous studies using mouse models of infection suggests that failure to establish a robust peripheral immune response to RVFV infection allows for viral spread to the brain, and exposure via the respiratory route increases the likelihood of bypassing such a protective response [4, 5, 26, 27, 50]. Thus, these studies suggest that the primary function of a protective response is to prevent the virus from ever reaching the brain. However, a recent study using a rat model of infection provided additional insight into the role of immune responses within the brain. Albe *et. al* [7] observed that rats infected subcutaneously with RVFV had detectable viral RNA in the brain as early as one day post infection despite lacking overt signs of disease. Furthermore, T cell infiltration into the brain was associated survival of infection, suggesting that functional T cell responses in the brain are ultimately capable of clearing infection and promoting survival and recovery. Rats exposed to aerosolized RVFV did not display T cell infiltration in the brain, and ultimately succumbed to infection. This study is consistent with the hypothesis that the lack of a robust peripheral immune response leads to increased pathogenesis within the brain, however it suggests that extension of peripheral responses to the brain is critical for the resolution of RVFV infection. Our present work indicates that innate immune responses to RVFV are robustly active within the brain. Thus, the role that cells within the brain play in recruiting and orchestrating a protective response warrants further investigation.

Microglia express a number of PRRs and respond to a wide variety of infectious agents in a diverse manner and can alternately establish an anti- or pro-inflammatory environment, depending on the specific genes that are induced as well as differing factors in the surrounding environment [51, 52]. Our results indicate that RVFV readily infects microglia, which then primarily utilize the RLR pathway via the signaling adaptor MAVS to detect and respond to RVFV infection. *In vitro*, microglia derived from *Mavs*^*-/-*^ animals displayed abrogated antiviral responses including diminished type I IFN and IFN-responsive gene expression and diminished capacity to upregulate surface markers of activation or secrete cytokines. Despite being severely attenuated, some response to infection was still noted, therefore we cannot rule out a minor role for other receptor(s) in the response of microglia to RVFV infection. Evaluation *in vivo* within *Tlr3*^*-/-*^ or *Tlr7*^*-/-*^ mice may reveal a role for these receptors, which might vary in importance within specific cell and tissue types.

The response of microglia to RVFV infection *in vivo* was also dependent on MAVS. Using scRNA-seq, we demonstrated that microglia in WT infected brains express type I IFNs and interferon responsive genes. Pathways that were enriched in WT microglia versus *Mavs*^*-/-*^ included response to virus and negative regulation of viral genome replication, indicating that WT microglia have a greater capacity to control viral infection. Specific ISGs with demonstrated antiviral activity were elevated including *Ifit1* and *Ifit3*, both of which have previously been shown to bind RVFV genomic RNA and have inhibitory activity on viral growth [53]. BST-2 has previously been shown to inhibit replication of several viruses, including HIV-1 and Lassa Virus, but was not capable of inhibiting RVFV replication in the cell types tested [54, 55]. Further investigation is required to determine if these ISGs have anti-RVFV activity in microglia. Other ISGs that were among the 20 most upregulated genes in WT microglia include master regulatory genes *Irf7* and *Isg15*, broad spectrum exonucluease *Isg20, Ifi27l2a*, which has been shown to have antiviral activity against WNV [56], and *Oasl1*, which can regulate *Irf7* expression [57]. Using flow cytometry, we demonstrated that WT microglia upregulate CD86 and I-A/I-E on the cell surface in response to RVFV infection. Upregulation of surface markers was restricted to microglia that were RVFV^+^, which were of relatively low abundance in WT brains. Separating microglia from infected brains into RVFV^+^ and RVFV^-^ populations was restricted to antibody staining and was not possible using scRNA-seq due to the sequencing technology being unable to capture RVFV genomic RNA. While we cannot ascertain which sequenced microglia were infected or uninfected, it can be inferred that the changes in gene expression observed within WT microglia were induced largely in trans in response to secreted factors from neighboring cells due to the low levels of infection detected by antibody staining.

In contrast, microglia from *Mavs*^*-/-*^ brains displayed much higher levels of infection, ranging from 15-50% of the total microglia. *Mavs*^*-/-*^ microglia did not express type I IFNs, and upregulated genes enriched in inflammatory processes such as response to interleukin-1 and positive regulation of interleukin-6. *Mavs*^*-/-*^ microglia upregulated surface expression of I-A/I-E, on both RVFV^+^ and RVFV^-^ microglia *in vivo*. Antigen processing and presentation was observed transcriptionally only within WT microglia, therefore the functional significance of I-A/I-E expression is unclear. Interestingly, *Irf1, Irf8*, and *Il1b* were amongst the genes most upregulated in *Mavs*^*-/-*^ microglia versus WT. Previous studies utilizing a mouse model of nerve injury indicated increased *Il1b* expression by microglia that was dependent on *Irf1* and *Irf8* expression [39, 40]. These studies in conjunction with our present study suggest that different types of pathological insults can activate a common inflammatory state in microglia and further suggest that failure to establish an appropriate antiviral response may dispose the microglia to remain in a proinflammatory state.

Our results highlight the importance of MAVS in orchestrating a protective immune response against RVFV infection in the brain. The lack of MAVS resulted in a dysregulated immune response, with more inflammatory gene expression and less functional adaptive immune responses. The brain infiltrating lymphocytes consisted of T and NK cells in both WT and *Mavs*^*-/-*^ animals however more NK than T cells were observed in WT. Furthermore, the T and NK cells in the brains of *Mavs*^*-/-*^ mice appear to be functionally deficient due to decreased expression of ISGs and genes related to killing functions. Thus, while more cells are recruited to the brain, likely due to the increased presence of viral antigen, they are less able to control viral replication. This is in line with previous studies investigating the role of MAVS in WNV infection which demonstrated increased infiltration of T cells with lower functional avidity into the brains of *Mavs*^*-/-*^ mice [30, 58]. Further work is needed to characterize the functional deficiency of T cells in *Mavs*^*-/-*^ mice during RVFV infection and to understand the molecular details that govern this deficiency. Moreover, the role of NK cells in RVFV infection has not been investigated, and our data suggests that NK cells may be actively recruited to the brain to help control RVFV infection. Interestingly, we observed increased expression of *l15, Il18*, and *Il27*, which have been shown to activate T and NK cells [46-49], in WT microglia and other myeloid cells. Secretion of these cytokines may be one mechanism by which microglia regulate the cytotoxic activity of lymphocytes, and lack of expression in *Mavs*^*-/-*^ microglia could be playing a role in the defective lymphocyte responses observed in these animals. Future studies will explore the relationship between secreted factors from microglia and the induction of protective T and NK cell responses during RVFV infection of the brain. Taken together, previous studies and our present study indicate that RLR signaling through MAVS is required not only to control viral replication early but is also necessary to induce a fully functional adaptive response.

In summary, this present work provides a better understanding of the immune response in the brain to RVFV infection. Furthermore, it defines a protective role for MAVS in propagating antiviral responses in the brain and suggests that signaling through MAVS may also be required for functional T and NK cell responses in the brain. Better understanding of the immune responses that are active against RVFV in the brain may contribute to therapeutics that effectively harness or augment these responses and lead to treatments for encephalitic disease.

## Materials and Methods

### Cells and viruses

Vero, EOC 13.31, SIM A9, and LADMAC cells were obtained from the American Type Culture Collection (ATCC). Cell lines were maintained in the following culture media: Vero: Dulbecco’s modified Eagle’s medium (DMEM) supplemented with 10% fetal bovine serum (FBS). EOC 13.31: DMEM supplemented with 10% FBS and 20% LADMAC conditioned media to provide a source of CSF-1 to support microglial growth [59]. SIM-A9: DMEM:F12 supplemented with 10% FBS and 5% horse serum. LADMAC: Minimal essential medium supplemented with 10% FBS. Primary glial cells: RPMI 1640 supplemented with 10% FBS and 0.01 mg/mL gentamicin and 0.25 μg/mL amphotericin. All media was supplemented with 100 units/ml penicillin and 100 μg/ml streptomycin, and all cells were maintained at 37 °C in 5% CO2. All cell culture reagents were obtained from Thermo Fisher. LADMAC conditioned media was prepared by collecting supernatant from confluent cells that have been in culture for 5-7 days and centrifuged at 300 x *g* for 10 min to remove cellular debris.

Wild type Rift Valley fever virus (RVFV) strain ZH-501 was obtained from the NIH Biodefense and Emerging Infections Research Resources Repository, NIAID, NIH. The MP-12 strain was kindly provided by Oscar Negrete (Sandia National Laboratory). RVFV stocks were propagated in Vero cells as previously described [60, 61]. Titers of viral stocks were determined by standard plaque assay consisting of an agarose overlay and crystal violet staining. All work with the ZH501 strain was performed in Institutional Biosafety Committee approved BSL-3 and ABSL-3 facilities at Lawrence Livermore National Laboratory using appropriate PPE and protective measures.

### Mice

All animal work was conducted in accordance with protocols approved by the Lawrence Livermore National Laboratory Institutional Animal Care and Use Committee. C57BL/6 as well as mice genetically deficient in MAVS (B6;129-*Mavs*^*tm1Zjc*^*/J*; Jax stock No: 008634), TLR3 B6;129S1-*Tlr3*^*tm1Flv*^/J; Jax Stock No: 005217), and TLR7 (B6.129S1-*Tlr7*^*tm1Flv*^/J; Jax stock No: 008380) were obtained from Jackson Laboratory. For experiments using knockout (KO) mice, animals were crossed to WT C57BL/6 mice to generate a heterozygous F1 generation. F1 littermates were crossed to generate homozygous WT and KO F2 progeny. Matched WT and KO animals from the same generation were used for each genotype. All animals were maintained in PHS-assured facilities.

### Isolation of primary microglia isolation

Primary microglia were isolated and cultured as described previously [62, 63]. Briefly, brains from 1 – 4-day old neonatal mice were dissected to remove meninges and large blood vessels and finely minced with sterile surgical scissors. The minced tissue was then forced through a 70 μM cell strainer (Fischer Scientific) and rinsed with cold glial media. Cells were pelleted at 300 x *g* for 10 min then resuspended in 20 mL fresh media and placed in a T75 flask (1 flask per 7 – 9 brains) and maintained in culture for 2 weeks. Microglia were harvested from the mixed glial culture by shaking flasks for 4 h at 200 rpm using an orbital shaker. Cells were pelleted at 300 x *g* for 10 min and resuspended in 10 mL microglia growth media (glial media + 20% LADMAC conditioned media). Microglia from up to 3 T75 flasks were combined and placed in a T25 flask. Microglia were maintained in culture for up to one week before use in viral infection experiments.

### *In vitro* infection of microglia

Microglia were plated in 24 well tissue culture treated plate at a density of 250,000 cells per well primary glial cell media. Cells were infected with RVFV at a MOI of 5 (qPCR analysis) or 2 (flow cytometry/cytokine analysis) in primary glial cell media. Cells were incubated with virus for 4 h at 37 °C in 5% CO2. Viral infection media was then removed, and cells used for qPCR analysis were lysed for RNA extraction. Cells used for flow cytometry and cytokine analysis were washed one time with PBS, then replenished with fresh media and incubated for another 18-24 h. Supernatants were then removed and stored at -80 °C for cytokine analysis, and cells were processed for flow cytometric analysis. Data for all *in vitro* assays is displayed as the average from 3 triplicate wells and is representative of an experiment performed at least twice.

### Cytokine analysis

Cytokines were quantified using Legendplex multiplex bead-based assay (Biolegend) using the mouse anti-virus response panel according to manufacturer’s instructions. Flow cytometry of the beads was performed using a FACSAria Fusion and data were analyzed using Biolegend’s cloud-based analysis software available at https://legendplex.qognit.com.

### *In vivo* infection

Groups of male and female *Mavs*^*-/-*^ and wildtype (WT) control littermates ranging in age from 8-12 weeks were inoculated intranasally with 5×10^5^ (MP-12, n=12) or 1000 PFU (ZH501, n=20) RVFV while under anesthesia (4-5% isoflurane in 100% oxygen). Mice were monitored daily for signs of morbidity and animals were humanely euthanized upon signs of severe disease by CO_2_ asphyxiation. For tissue harvest, animals were euthanized by CO_2_ asphyxiation and the whole animal was perfused with 30 mL sterile PBS containing 50,000 U/L sodium heparin *via* the left ventricle.

### Brain tissue isolation and preparation

Preparation of brain tissue for flow cytometric, RNA sequencing, cytokine, and viral titer analysis was performed as previously described [64]. Following euthanasia and perfusion, brains were removed and placed in digestion buffer (PBS pH 7.4 (Thermo Fisher) + liberase + DNase I (both from Roche) to a final concentration of 1.6 wunsch/mL and 0.5 mg/mL, respectively) on ice in a 1.5 mL tube. Brains were finely diced into 1-2 mm^3^ pieces with small scissors. 3-4 pieces were placed into RNAlater (Qiagen) for gene expression analysis. The remaining tissue was digested at 37 °C for 30 min. EDTA was added to a final concentration of 10 mM to stop the digestion reaction. A cell suspension was generated by gentle pipetting followed by passage through a 70 μm cell strainer. The cell strainer was rinsed with PBS supplemented with 5% FBS to a total volume of 20 mL. Aliquots of this suspension were stored at -80 °C for cytokine and viral titer analysis. The remaining suspension was subjected to Percoll gradient centrifugation to purify mononuclear immune cells for flow cytometric analysis as previously described [64].

### Flow cytometry

Cells were incubated for 30 min on ice in 100 μl Hank’s balanced salt solution (Thermo Fisher) + 2% FBS with Fc block (1:100 dilution, clone 2.4G2; BD Biosciences) along with the following antibodies (all from BD Biosciences): CD45 APC-Cy7 (1:500, clone 30-F11), CD11b AF488 (1:500, clone M1/70), CD80 BV421 (1:200, clone 16-1OA1), and CD86 PE-Cy7 (1:500, clone GL1). IA/I-E AF647 (1:500, clone M5/114.15.2; BioLegend). Cells were then fixed and permeabilized using BD Cytofix/Cytoperm (BD Biosciences) according to manufacturer’s instructions. Cells were then incubated with an anti-RVFV antibody (kindly provided by Dr. Robert Tesh and the World Reference Center of Emerging Viruses and Arboviruses) at 1:500 dilution, followed by goat anti-mouse PE secondary antibody (1:1000, Santa Cruz biotechnology). Flow cytometry was performed using a FACSAria Fusion and data were analyzed using FlowJo software. Microglia, other myeloid lineage, and lymphocytes were resolved using CD45 and CD11b expression, with microglia identified as CD45^int^ CD11b^int^, other myeloid as CD45^hi^ CD11b^hi^, and lymphocytes as CD45^hi^ CD11b^−^ as previously described [64].

### qPCR and bulk RNA seq analyses

RVFV genomic RNA was quantified in brain tissue as described previously [5, 26, 65]. To normalize between samples, results were normalized to GAPDH expression. For *in vitro* qPCR analysis of immune gene expression, RNA from infected and uninfected microglia was harvested at 4 h post infection using RNeasy plus kits, and cDNA was generated using RT^2^ First Strand Synthesis kit (both from Qiagen) according to manufacturer’s instructions. Real-time quantitative RT-PCR analysis of the samples was carried out using the mouse antiviral response RT^2^ Profiler PCR array (Qiagen) on a 7900HT Fast Real-Time PCR system (Thermo Fisher) according to manufacturer’s instructions. Data were analyzed using Qiagen’s online analysis software available at https://geneglobe.qiagen.com/us/analyze. Data are shown as the log_2_ fold change in gene expression in infected versus uninfected samples.

For bulk sequencing analysis, RNA was harvested from RNAlater preserved brain tissue using RNeasy Plus kit. Poly(A)+-enriched cDNA libraries were generated using the Illumina TruSeq RNA Library Prep kit v2 (Illumina Inc). The 75 bp single-end reads were sequenced was performed using an Illumina (Illumina Inc) NextSeq 500 instrument. Sequencing data quality was checked using FastQC software. Reads were mapped to the mouse reference genome (mm10) using STAR [66]. Read counts per gene locus were summarized with featureCounts [67]. Then the data was normalized using RUVseq [68] to correct for batch effects and other unwanted variations. Genes differentially expressed between uninfected and infected samples were identified using edgeR [69]. Gene ontology (GO) and pathway enrichment analysis was performed using functional annotation tool ToppGene [70]. Heatmaps were generated using heatmap.2 function in ‘gplots’ R package.

### Single cell RNA sequencing

Mice were euthanized and perfused as described above. Isolated brain tissues were immediately processed using a modified protocol from [71]. Cortices were dissected in cold Hibernate A medium (BrainBits LLC) and sliced to approximately 0.5 mm before transferred to a 15 mL falcon tube with Hibernate A and B27 medium (HABG) (Thermo Fischer Scientific). Collected tissue samples in 15 mL conical tubes were warmed up to 30°C in a shaking water bath for 8 minutes, before the HABG supernatant replaced with activated papain (34 U/mL, Worthington Biochemical Corporation) and the tubes placed back into the shaking water bath for tissue digestion (30°C, 150 rpm) for 30 minutes. Cells were released from the digested tissues by trituration using a fire-polished Pasteur pipette. Released cells were collected in the supernatant and filtered through a 70 mm MACS Smart Strainer (Miltenyi Biotec) into a new 15 mL conical tube. The single cell suspension was layered onto a Optiprep density gradient to separate cells from debris, after centrifugation (800 x *g*, 15 min, 22°C). The debris fraction was collected, and the gradient material diluted with HAGB before tubes were centrifuged (200 x *g*, 5 min, 22°C). The supernatant was aspirated, and ACK lysis buffer (Thermo Fisher Scientific) was added to the cell suspension remove any remaining red blood cells (5 min, RT). Hank’s balanced salt solution (Thermo Fisher) was added to the cell suspension containing the lysis buffer and tubes were centrifuged (200 x *g*, 5 min, 22°C). To remove dead cells from the single cell suspension, a Dead cell removal kit (Miltenyi Biotec) was used as directed by the vendor.

Cell pellets were resuspended in PBS with 0.04% non-acetylated BSA and counted on a Countess II automated cell counter prior to single-cell sequencing preparation using Chromium Single-cell 3’ GEM, Library & Gel Bead Kit v3 (10x Genomics Cat # 1000075) on a 10× Genomics Chromium Controller following manufacturers’ protocol. Sequencing data was demultiplexed, quality controlled, and analyzed using Cell Ranger (10x Genomics) and Seurat [72]. The Cell Ranger Single-Cell Software Suite was used to perform sample demultiplexing, barcode processing, and single-cell 3′gene counting. Samples were first demultiplexed and then aligned to the mouse genome (mm10) using “cellranger mkfastq” with default parameters. Unique molecular identifier counts were generated using “cellranger count”. Further analysis was performed using Seurat [72]. First, cells with fewer than 500 detected genes per cell and genes that were expressed by fewer than 5 cells were filtered out. After pre-processing, we performed data normalization, scaling, and identified 2000 most variable features. Then, anchors for data integration were identified using the ‘Find-IntegrationAnchors’ function. Next, these anchors were passed to the ‘IntegrateData’ function and new integrated matrix with all four datasets were generated. Subsequently, dimensionality reduction, clustering, and visualization were performed in Seurat as described before [73]. Genes differentially expressed between clusters were identified using ‘FindMarkers’ function implemented in Seurat.

## Acknowledgements and Funding

Funding for this research was provided by internal LLNL Laboratory Directed Research and Development funds (17-LW-038 and 22-ERD-038 to D.W.). The funders had no role in study design, data collection and analysis, decision to publish, or preparation of the manuscript. This work was performed under the auspices of the U.S. Department of Energy by Lawrence Livermore National Security, LLC, Lawrence Livermore National Laboratory under Contract DE-AC52-07NA27344.

## Author Contributions Statement

D.W., N.H., and F.B. conceived and designed the experiments. D.W., F.B., N.H., A.P., D.L., K.S., and A.R. conducted all *in vitro* and *in vivo* experiments described and D.W., N.H., F.B., and A.S. performed the data analysis. D.W., N.H., and G.L. wrote the manuscript. All authors contributed to editing the manuscript and approved the final version.

---

**Figure S1.**
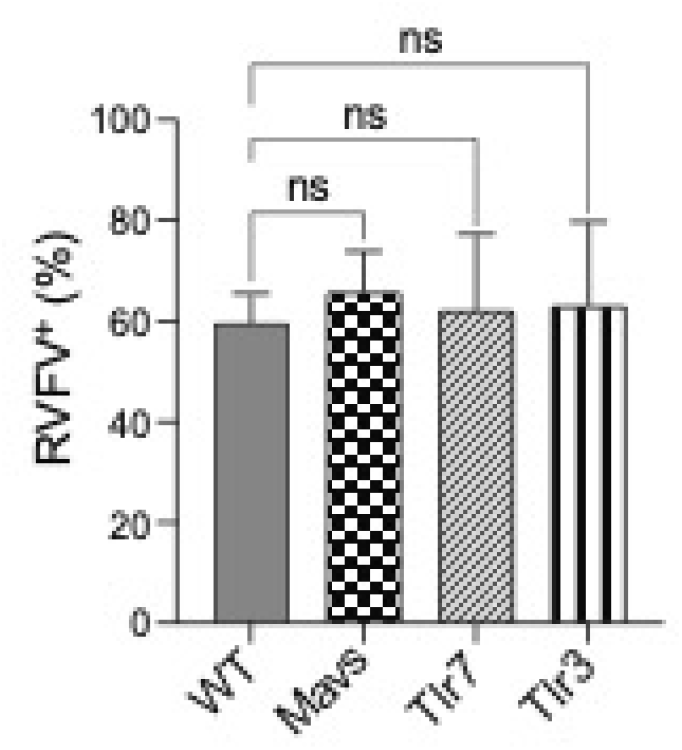
Infection of primary microglia is not dependent upon genotype. Primary microglia derived from WT or the indicated genetically deficient mice were infected with RVFV MP-12 and the percentage of cells positive for RVFV was assessed by flow cytometry.

**Figure S2.**
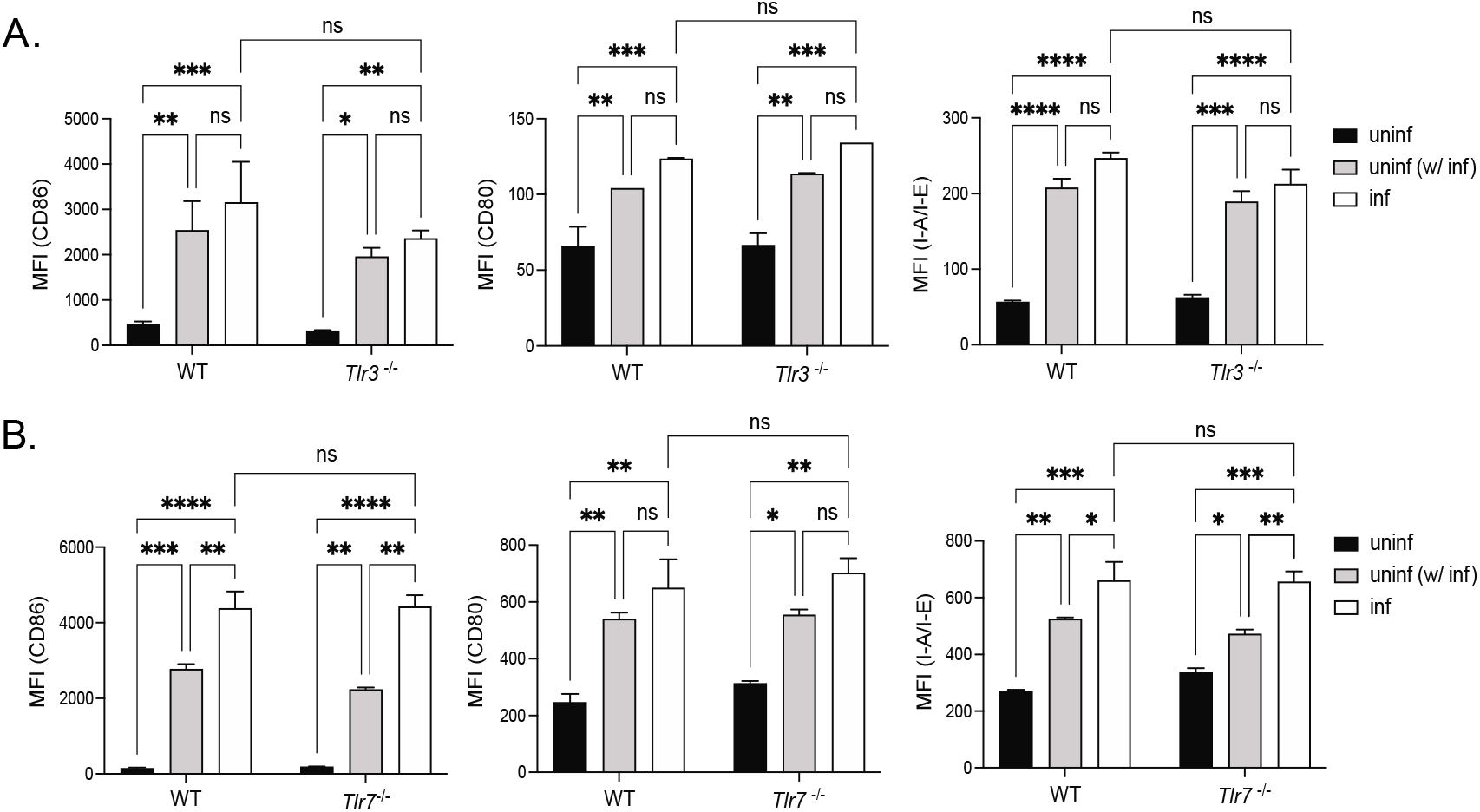
Expression of activation markers is not dependent on TLR3 or TLR7. Microglia derived from WT or *Tlr3*^*-/-*^ (A) or *Tlr7*^*-/-*^ (B) mice were infected with RVFV MP-12 and at 18-24 hours post-infection, cells were harvested for flow cytometry. The expression levels of the indicated activation markers were assessed on uninfected cells (black bars), uninfected cells in culture with infected cells (uninf (w/ inf), gray bars), and infected cells (inf, white bars) and shown as the mean fluorescence intensity of the indicated activation markers. Data are shown as the mean +/- SD *p <0.05, **p<0.01, ***p<0.001, ****p<0.0001

**Figure S3.**
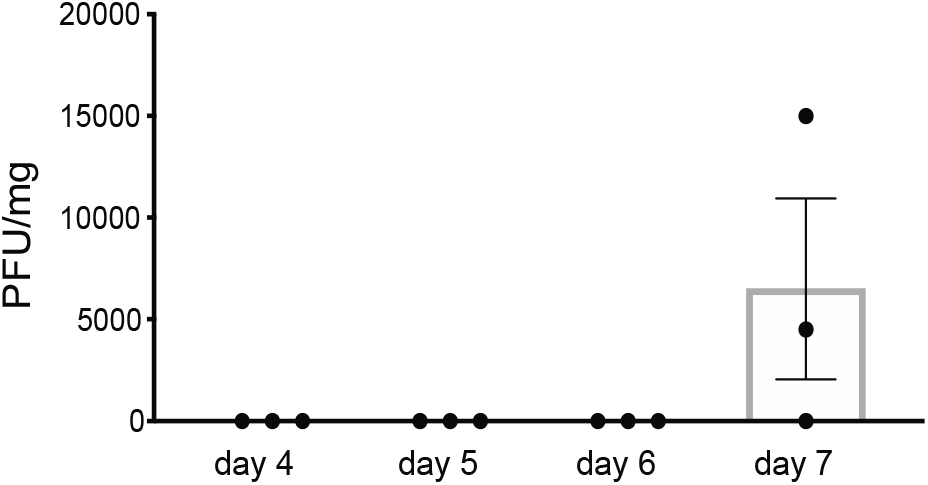
RVFV is detected in the brain on day 7 post infection. WT mice were infected intranasally with 5×10^5^ PFU RVFV MP-12 and brains were harvested on the indicated day post infection for viral quantitation.

**Figure S4.**
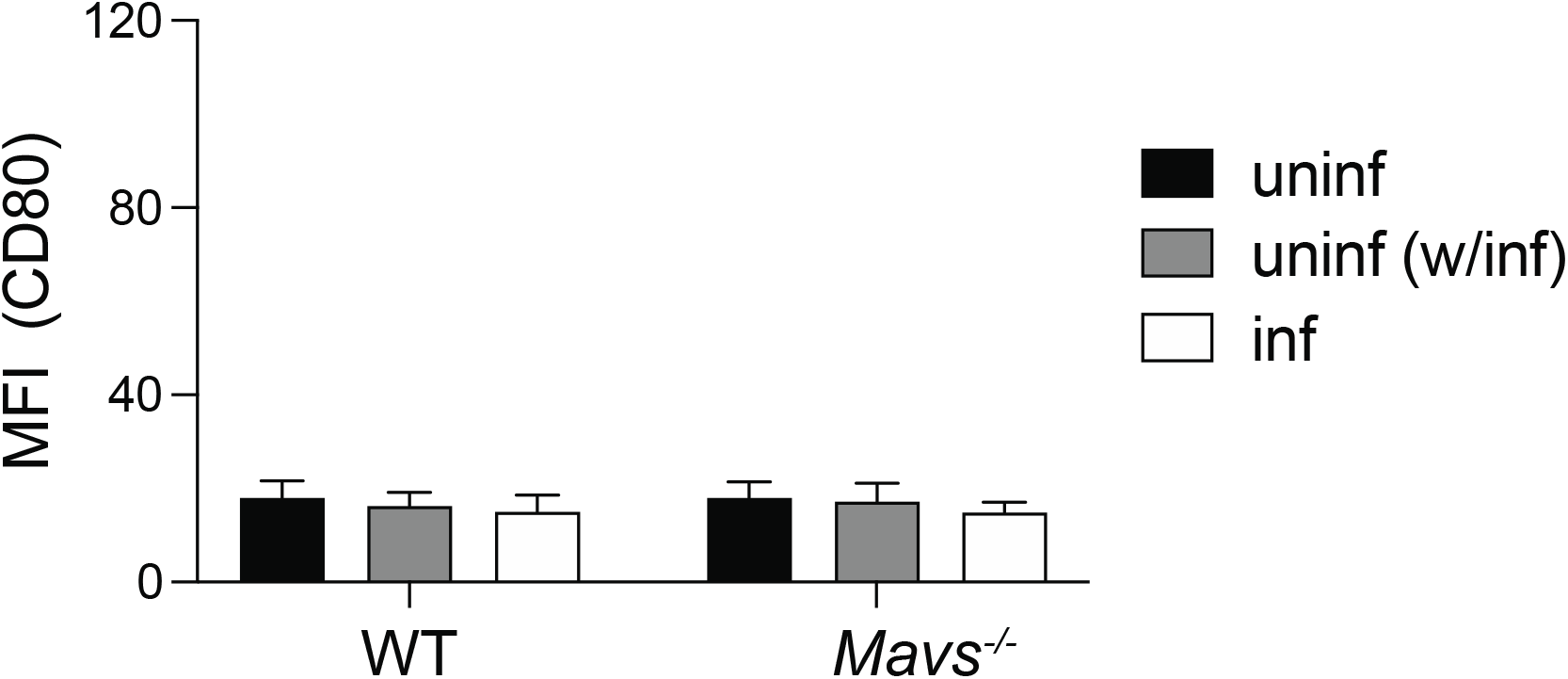
CD80 expression on microglia. Mean fluorescence intensity of CD80 on microglia isolated from the brains of WT and *Mavs*^*-/-*^ mice, +/- RVFV infection.

**Figure S5.**
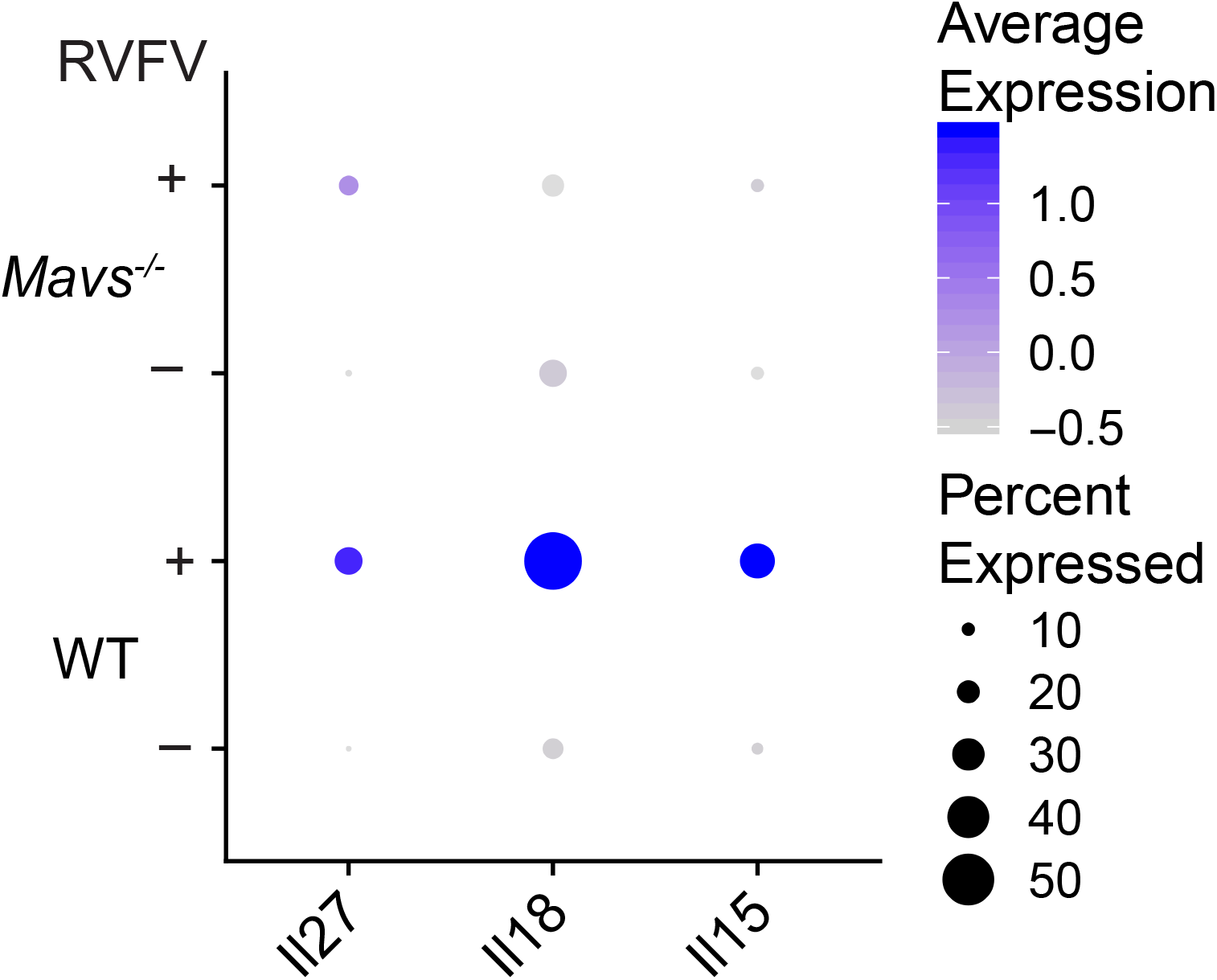
Microglia and myeloid cells from *Mavs*^*-/-*^ infected brains display lower levels of cytokine gene expression.

## References

1. Pepin M, Bouloy M, Bird BH, Kemp A, Paweska J. Rift Valley fever virus(Bunyaviridae: Phlebovirus): an update on pathogenesis, molecular epidemiology, vectors, diagnostics and prevention. Vet Res. 2010;41(6):61. doi: 10.1051/vetres/2010033. PubMed PMID: 21188836; PubMed Central PMCID: PMCPMC2896810.

2. Rolin AI, Berrang-Ford L, Kulkarni MA. The risk of Rift Valley fever virus introduction and establishment in the United States and European Union. Emerg Microbes Infect. 2013;2:e81. doi: 10.1038/emi.2013.81.

3. https://www.selectagents.gov/sat/list.htm.

4. Reed C, Lin K, Wilhelmsen C, Friedrich B, Nalca A, Keeney A, et al. Aerosol Exposure to Rift Valley Fever Virus Causes Earlier and More Severe Neuropathology in the Murine Model, which Has Important Implications for Therapeutic Development. PLOS Neglected Tropical Diseases. 2013;7(4):e2156. doi: 10.1371/journal.pntd.0002156.

5. Dodd KA, McElroy AK, Jones TL, Zaki SR, Nichol ST, Spiropoulou CF. Rift valley Fever virus encephalitis is associated with an ineffective systemic immune response and activated T cell infiltration into the CNS in an immunocompetent mouse model. PLoS Negl Trop Dis. 2014;8(6):e2874. Epub 2014/06/13. doi: 10.1371/journal.pntd.0002874. PubMed PMID: 24922480; PubMed Central PMCID: PMCPMC4055548.

6. Ermler ME, Yerukhim E, Schriewer J, Schattgen S, Traylor Z, Wespiser AR, et al. RNA Helicase Signaling Is Critical for Type I Interferon Production and Protection against Rift Valley Fever Virus during Mucosal Challenge. Journal of Virology. 2013;87(9):4846–60. doi: 10.1128/jvi.01997-12.

7. Albe JR, Boyles DA, Walters AW, Kujawa MR, McMillen CM, Reed DS, et al. Neutrophil and macrophage influx into the central nervous system are inflammatory components of lethal Rift Valley fever encephalitis in rats. PLoS Pathog. 2019;15(6):e1007833. Epub 2019/06/21. doi: 10.1371/journal.ppat.1007833. PubMed PMID: 31220182; PubMed Central PMCID: PMCPMC6605717.

8. Anyangu AS, Gould LH, Sharif SK, Nguku PM, Omolo JO, Mutonga D, et al. Risk Factors for Severe Rift Valley Fever Infection in Kenya, 2007. The American Journal of Tropical Medicine and Hygiene. 2010;83(2 Suppl):14–21. doi: 10.4269/ajtmh.2010.09-0293.

9. LaBeaud AD, Pfeil S, Muiruri S, Dahir S, Sutherland LJ, Traylor Z, et al. Factors Associated with Severe Human Rift Valley Fever in Sangailu, Garissa County, Kenya. PLoS Negl Trop Dis. 2015;9(3):e0003548. doi: 10.1371/journal.pntd.0003548.

10. Madani TA, Al-Mazrou YY, Al-Jeffri MH, Mishkhas AA, Al-Rabeah AM, Turkistani AM, et al. Rift Valley fever epidemic in Saudi Arabia: epidemiological, clinical, and laboratory characteristics. Clin Infect Dis. 2003;37(8):1084–92. Epub 2003/10/03. doi: 10.1086/378747. PubMed PMID: 14523773.

11. Javelle E, Lesueur A, Pommier de Santi V, de Laval F, Lefebvre T, Holweck G, et al. The challenging management of Rift Valley Fever in humans: literature review of the clinical disease and algorithm proposal. Ann Clin Microbiol Antimicrob. 2020;19(1):4-. doi: 10.1186/s12941-020-0346-5. PubMed PMID: 31969141.

12. Rustenhoven J. A privileged brain. Science. 2021;374(6567):548. Epub 20211028. doi: 10.1126/science.abl7122. PubMed PMID: 34709902.

13. Lampron A, Elali A, Rivest S. Innate immunity in the CNS: redefining the relationship between the CNS and Its environment. Neuron. 2013;78(2):214–32. Epub 2013/04/30. doi: 10.1016/j.neuron.2013.04.005. PubMed PMID: 23622060.

14. Carty M, Reinert L, Paludan SR, Bowie AG. Innate antiviral signalling in the central nervous system. Trends Immunol. 2014;35(2):79–87. Epub 20131206. doi: 10.1016/j.it.2013.10.012. PubMed PMID: 24316012.

15. Singh H, Koury J, Kaul M. Innate Immune Sensing of Viruses and Its Consequences for the Central Nervous System. Viruses. 2021;13(2). Epub 2021/01/28. doi: 10.3390/v13020170. PubMed PMID: 33498715; PubMed Central PMCID: PMCPMC7912342.

16. Denizot M, Neal JW, Gasque P. Encephalitis due to emerging viruses: CNS innate immunity and potential therapeutic targets. The Journal of infection. 2012;65(1):1–16. Epub 2012/04/10. doi: 10.1016/j.jinf.2012.03.019. PubMed PMID: 22484271.

17. Drokhlyansky E, Göz Aytürk D, Soh TK, Chrenek R, O’Loughlin E, Madore C, et al. The brain parenchyma has a type I interferon response that can limit virus spread. Proc Natl Acad Sci U S A. 2017;114(1):E95–e104. Epub 2016/12/17. doi: 10.1073/pnas.1618157114. PubMed PMID: 27980033; PubMed Central PMCID: PMCPMC5224383.

18. Moseman EA, Blanchard AC, Nayak D, McGavern DB. T cell engagement of cross-presenting microglia protects the brain from a nasal virus infection. Sci Immunol. 2020;5(48). Epub 2020/06/07. doi: 10.1126/sciimmunol.abb1817. PubMed PMID: 32503876; PubMed Central PMCID: PMCPMC7416530.

19. Sanchez JMS, DePaula-Silva AB, Doty DJ, Hanak TJ, Truong A, Libbey JE, et al. The CSF1R-Microglia Axis Has Protective Host-Specific Roles During Neurotropic Picornavirus Infection. Frontiers in Immunology. 2021;12(3568). doi: 10.3389/fimmu.2021.621090.

20. Funk KE, Klein RS. CSF1R antagonism limits local restimulation of antiviral CD8+ T cells during viral encephalitis. Journal of neuroinflammation. 2019;16(1):22. doi: 10.1186/s12974-019-1397-4.

21. Mangale V, Syage AR, Ekiz HA, Skinner DD, Cheng Y, Stone CL, et al. Microglia influence host defense, disease, and repair following murine coronavirus infection of the central nervous system. Glia. 2020;68(11):2345–60. Epub 20200525. doi: 10.1002/glia.23844. PubMed PMID: 32449994; PubMed Central PMCID: PMCPMC7280614.

22. Chhatbar C, Detje CN, Grabski E, Borst K, Spanier J, Ghita L, et al. Type I Interferon Receptor Signaling of Neurons and Astrocytes Regulates Microglia Activation during Viral Encephalitis. Cell Rep. 2018;25(1):118-29.e4. Epub 2018/10/04. doi: 10.1016/j.celrep.2018.09.003. PubMed PMID: 30282022; PubMed Central PMCID: PMCPMC7103936.

23. Tsai TT, Chen CL, Lin YS, Chang CP, Tsai CC, Cheng YL, et al. Microglia retard dengue virus-induced acute viral encephalitis. Sci Rep. 2016;6:27670. Epub 20160609. doi: 10.1038/srep27670. PubMed PMID: 27279150; PubMed Central PMCID: PMCPMC4899773.

24. Tansey MG, Goldberg MS. Neuroinflammation in Parkinson’s disease: its role in neuronal death and implications for therapeutic intervention. Neurobiol Dis. 2010;37(3):510–8. Epub 20091110. doi: 10.1016/j.nbd.2009.11.004. PubMed PMID: 19913097; PubMed Central PMCID: PMCPMC2823829.

25. Zhang G, Wang Z, Hu H, Zhao M, Sun L. Microglia in Alzheimer’s Disease: A Target for Therapeutic Intervention. Front Cell Neurosci. 2021;15:749587. Epub 20211124. doi: 10.3389/fncel.2021.749587. PubMed PMID: 34899188; PubMed Central PMCID: PMCPMC8651709.

26. Dodd KA, McElroy AK, Jones ME, Nichol ST, Spiropoulou CF. Rift Valley fever virus clearance and protection from neurologic disease are dependent on CD4+ T cell and virus-specific antibody responses. J Virol. 2013;87(11):6161–71. Epub 20130327. doi: 10.1128/jvi.00337-13. PubMed PMID: 23536675; PubMed Central PMCID: PMCPMC3648110.

27. Harmon JR, Spengler JR, Coleman-McCray JD, Nichol ST, Spiropoulou CF, McElroy AK. CD4 T Cells, CD8 T Cells, and Monocytes Coordinate To Prevent Rift Valley Fever Virus Encephalitis. J Virol. 2018;92(24). Epub 20181127. doi: 10.1128/jvi.01270-18. PubMed PMID: 30258000; PubMed Central PMCID: PMCPMC6258944.

28. Daffis S, Samuel MA, Suthar MS, Gale M, Jr., Diamond MS. Toll-like receptor 3 has a protective role against West Nile virus infection. J Virol. 2008;82(21):10349–58. Epub 2008/08/22. doi: 10.1128/jvi.00935-08. PubMed PMID: 18715906; PubMed Central PMCID: PMCPMC2573187.

29. Town T, Bai F, Wang T, Kaplan AT, Qian F, Montgomery RR, et al. Toll-like receptor 7 mitigates lethal West Nile encephalitis via interleukin 23-dependent immune cell infiltration and homing. Immunity. 2009;30(2):242–53. Epub 2009/02/10. doi: 10.1016/j.immuni.2008.11.012. PubMed PMID: 19200759; PubMed Central PMCID: PMCPMC2707901.

30. Suthar MS, Ma DY, Thomas S, Lund JM, Zhang N, Daffis S, et al. IPS-1 is essential for the control of West Nile virus infection and immunity. PLoS Pathog. 2010;6(2):e1000757. Epub 2010/02/09. doi: 10.1371/journal.ppat.1000757. PubMed PMID: 20140199; PubMed Central PMCID: PMCPMC2816698.

31. Hise AG, Traylor Z, Hall NB, Sutherland LJ, Dahir S, Ermler ME, et al. Association of Symptoms and Severity of Rift Valley Fever with Genetic Polymorphisms in Human Innate Immune Pathways. PLoS Negl Trop Dis. 2015;9(3):e0003584. doi: 10.1371/journal.pntd.0003584.

32. Roberts KK, Hill TE, Davis MN, Holbrook MR, Freiberg AN. Cytokine response in mouse bone marrow derived macrophages after infection with pathogenic and non-pathogenic Rift Valley fever virus. J Gen Virol. 2015;96(Pt 7):1651–63. Epub 2015/03/12. doi: 10.1099/vir.0.000119. PubMed PMID: 25759029; PubMed Central PMCID: PMCPMC4635452.

33. Artegiani B, Lyubimova A, Muraro M, van Es JH, van Oudenaarden A, Clevers H. A Single-Cell RNA Sequencing Study Reveals Cellular and Molecular Dynamics of the Hippocampal Neurogenic Niche. Cell Rep. 2017;21(11):3271–84. doi: 10.1016/j.celrep.2017.11.050. PubMed PMID: 29241552.

34. Cougnoux A, Yerger JC, Fellmeth M, Serra-Vinardell J, Martin K, Navid F, et al. Single Cell Transcriptome Analysis of Niemann-Pick Disease, Type C1 Cerebella. Int J Mol Sci. 2020;21(15). Epub 20200728. doi: 10.3390/ijms21155368. PubMed PMID: 32731618; PubMed Central PMCID: PMCPMC7432835.

35. Vanlandewijck M, He L, Mäe MA, Andrae J, Ando K, Del Gaudio F, et al. A molecular atlas of cell types and zonation in the brain vasculature. Nature. 2018;554(7693):475–80. Epub 20180214. doi: 10.1038/nature25739. PubMed PMID: 29443965.

36. Hammond TR, Dufort C, Dissing-Olesen L, Giera S, Young A, Wysoker A, et al. Single-Cell RNA Sequencing of Microglia throughout the Mouse Lifespan and in the Injured Brain Reveals Complex Cell-State Changes. Immunity. 2019;50(1):253-71.e6. Epub 20181121. doi: 10.1016/j.immuni.2018.11.004. PubMed PMID: 30471926; PubMed Central PMCID: PMCPMC6655561.

37. Morimoto K, Nakajima K. Role of the Immune System in the Development of the Central Nervous System. Frontiers in Neuroscience. 2019;13(916). doi: 10.3389/fnins.2019.00916.

38. Goldmann T, Wieghofer P, Jordão MJ, Prutek F, Hagemeyer N, Frenzel K, et al. Origin, fate and dynamics of macrophages at central nervous system interfaces. Nat Immunol. 2016;17(7):797–805. Epub 2016/05/03. doi: 10.1038/ni.3423. PubMed PMID: 27135602; PubMed Central PMCID: PMCPMC4968048.

39. Masuda T, Tsuda M, Yoshinaga R, Tozaki-Saitoh H, Ozato K, Tamura T, et al. IRF8 is a critical transcription factor for transforming microglia into a reactive phenotype. Cell Rep. 2012;1(4):334–40. Epub 2012/07/27. doi: 10.1016/j.celrep.2012.02.014. PubMed PMID: 22832225; PubMed Central PMCID: PMCPMC4158926.

40. Masuda T, Iwamoto S, Mikuriya S, Tozaki-Saitoh H, Tamura T, Tsuda M, et al. Transcription factor IRF1 is responsible for IRF8-mediated IL-1β expression in reactive microglia. J Pharmacol Sci. 2015;128(4):216–20. Epub 2015/09/01. doi: 10.1016/j.jphs.2015.08.002. PubMed PMID: 26318672.

41. Wieronska JM, Cieslik P, Kalinowski L. Nitric Oxide-Dependent Pathways as Critical Factors in the Consequences and Recovery after Brain Ischemic Hypoxia. Biomolecules. 2021;11(8). Epub 20210726. doi: 10.3390/biom11081097. PubMed PMID: 34439764; PubMed Central PMCID: PMCPMC8392725.

42. Voskoboinik I, Whisstock JC, Trapani JA. Perforin and granzymes: function, dysfunction and human pathology. Nature reviews Immunology. 2015;15(6):388–400. doi: 10.1038/nri3839. PubMed PMID: 25998963.

43. Shrestha B, Samuel MA, Diamond MS. CD8+ T cells require perforin to clear West Nile virus from infected neurons. J Virol. 2006;80(1):119–29. doi: 10.1128/jvi.80.1.119-129.2006. PubMed PMID: 16352536; PubMed Central PMCID: PMCPMC1317548.

44. Shi F-D, Ransohoff RM. Nature killer cells in the central nervous system. Natural Killer Cells. 2010:373–83. Epub 2010/01/29. doi: 10.1016/B978-0-12-370454-2.00028-4. PubMed PMID: PMC7150147.

45. Yao Y, Strauss-Albee DM, Zhou JQ, Malawista A, Garcia MN, Murray KO, et al. The natural killer cell response to West Nile virus in young and old individuals with or without a prior history of infection. PLOS ONE. 2017;12(2):e0172625. doi: 10.1371/journal.pone.0172625.

46. Choi YH, Lim EJ, Kim SW, Moon YW, Park KS, An HJ. IL-27 enhances IL-15/IL-18-mediated activation of human natural killer cells. J Immunother Cancer. 2019;7(1):168. Epub 20190705. doi: 10.1186/s40425-019-0652-7. PubMed PMID: 31277710; PubMed Central PMCID: PMCPMC6612093.

47. Ochayon DE, Waggoner SN. The Effect of Unconventional Cytokine Combinations on NK-Cell Responses to Viral Infection. Frontiers in Immunology. 2021;12(775). doi: 10.3389/fimmu.2021.645850.

48. Nakanishi K. Unique Action of Interleukin-18 on T Cells and Other Immune Cells. Frontiers in Immunology. 2018;9(763). doi: 10.3389/fimmu.2018.00763.

49. Ziblat A, Domaica CI, Spallanzani RG, Iraolagoitia XL, Rossi LE, Avila DE, et al. IL-27 stimulates human NK-cell effector functions and primes NK cells for IL-18 responsiveness. European journal of immunology. 2015;45(1):192–202. Epub 20141202. doi: 10.1002/eji.201444699. PubMed PMID: 25308526.

50. Smith DR, Steele KE, Shamblin J, Honko A, Johnson J, Reed C, et al. The pathogenesis of Rift Valley fever virus in the mouse model. Virology. 2010;407(2):256–67. Epub 2010/09/21. doi: 10.1016/j.virol.2010.08.016. PubMed PMID: 20850165.

51. Chen Z, Zhong D, Li G. The role of microglia in viral encephalitis: a review. Journal of neuroinflammation. 2019;16(1):76. Epub 2019/04/11. doi: 10.1186/s12974-019-1443-2. PubMed PMID: 30967139; PubMed Central PMCID: PMCPMC6454758.

52. Stonedahl S, Clarke P, Tyler KL. The Role of Microglia during West Nile Virus Infection of the Central Nervous System. Vaccines (Basel). 2020;8(3). Epub 2020/09/03. doi: 10.3390/vaccines8030485. PubMed PMID: 32872152; PubMed Central PMCID: PMCPMC7563127.

53. Pichlmair A, Lassnig C, Eberle CA, Górna MW, Baumann CL, Burkard TR, et al. IFIT1 is an antiviral protein that recognizes 5’-triphosphate RNA. Nat Immunol. 2011;12(7):624–30. Epub 2011/06/07. doi: 10.1038/ni.2048. PubMed PMID: 21642987.

54. Neil SJ, Zang T, Bieniasz PD. Tetherin inhibits retrovirus release and is antagonized by HIV-1 Vpu. Nature. 2008;451(7177):425–30. Epub 2008/01/18. doi: 10.1038/nature06553. PubMed PMID: 18200009.

55. Radoshitzky SR, Dong L, Chi X, Clester JC, Retterer C, Spurgers K, et al. Infectious Lassa virus, but not filoviruses, is restricted by BST-2/tetherin. J Virol. 2010;84(20):10569–80. Epub 2010/08/06. doi: 10.1128/jvi.00103-10. PubMed PMID: 20686043; PubMed Central PMCID: PMCPMC2950602.

56. Lucas TM, Richner JM, Diamond MS, Perlman S. The Interferon-Stimulated Gene <i>Ifi27l2a</i> Restricts West Nile Virus Infection and Pathogenesis in a Cell-Type- and Region-Specific Manner. Journal of Virology. 2016;90(5):2600–15. doi: doi:10.1128/JVI.02463-15.

57. Lee MS, Kim B, Oh GT, Kim Y-J. OASL1 inhibits translation of the type I interferon– regulating transcription factor IRF7. Nature Immunology. 2013;14(4):346–55. doi: 10.1038/ni.2535.

58. Zhao J, Vijay R, Zhao J, Gale M, Jr., Diamond MS, Perlman S. MAVS Expressed by Hematopoietic Cells Is Critical for Control of West Nile Virus Infection and Pathogenesis. J Virol. 2016;90(16):7098–108. Epub 20160727. doi: 10.1128/jvi.00707-16. PubMed PMID: 27226371; PubMed Central PMCID: PMCPMC4984631.

59. O’Keefe GM, Nguyen VT, Ping Tang L, Benveniste EN. IFN-γ Regulation of Class II Transactivator Promoter IV in Macrophages and Microglia: Involvement of the Suppressors of Cytokine Signaling-1 Protein. The Journal of Immunology. 2001;166(4):2260–9. doi: 10.4049/jimmunol.166.4.2260.

60. Harmon B, Schudel BR, Maar D, Kozina C, Ikegami T, Tseng CT, et al. Rift Valley fever virus strain MP-12 enters mammalian host cells via caveola-mediated endocytosis. J Virol. 2012;86(23):12954–70. Epub 20120919. doi: 10.1128/jvi.02242-12. PubMed PMID: 22993156; PubMed Central PMCID: PMCPMC3497621.

61. Harmon B, Bird SW, Schudel BR, Hatch AV, Rasley A, Negrete OA. A Genome-Wide RNA Interference Screen Identifies a Role for Wnt/β-Catenin Signaling during Rift Valley Fever Virus Infection. J Virol. 2016;90(16):7084–97. Epub 20160727. doi: 10.1128/jvi.00543-16. PubMed PMID: 27226375; PubMed Central PMCID: PMCPMC4984662.

62. Rasley A, Anguita J, Marriott I. Borrelia burgdorferi induces inflammatory mediator production by murine microglia. Journal of neuroimmunology. 2002;130(1):22–31. doi: https://doi.org/10.1016/S0165-5728(02)00187-X.

63. Rasley A, Bost KL, Olson JK, Miller SD, Marriott I. Expression of functional NK-1 receptors in murine microglia. Glia. 2002;37(3):258–67. doi: https://doi.org/10.1002/glia.10034.

64. Martin E, El-Behi M, Fontaine B, Delarasse C. Analysis of Microglia and Monocyte-derived Macrophages from the Central Nervous System by Flow Cytometry. Journal of visualized experiments : JoVE. 2017;(124):55781. doi: 10.3791/55781. PubMed PMID: 28671658.

65. Bird BH, Bawiec DA, Ksiazek TG, Shoemaker TR, Nichol ST. Highly sensitive and broadly reactive quantitative reverse transcription-PCR assay for high-throughput detection of Rift Valley fever virus. J Clin Microbiol. 2007;45(11):3506–13. Epub 2007/09/07. doi: 10.1128/jcm.00936-07. PubMed PMID: 17804663; PubMed Central PMCID: PMCPMC2168471.

66. Dobin A, Davis CA, Schlesinger F, Drenkow J, Zaleski C, Jha S, et al. STAR: ultrafast universal RNA-seq aligner. Bioinformatics. 2013;29(1):15–21. Epub 20121025. doi: 10.1093/bioinformatics/bts635. PubMed PMID: 23104886; PubMed Central PMCID: PMCPMC3530905.

67. Liao Y, Smyth GK, Shi W. featureCounts: an efficient general purpose program for assigning sequence reads to genomic features. Bioinformatics. 2014;30(7):923–30. Epub 20131113. doi: 10.1093/bioinformatics/btt656. PubMed PMID: 24227677.

68. Risso D, Ngai J, Speed TP, Dudoit S. Normalization of RNA-seq data using factor analysis of control genes or samples. Nature biotechnology. 2014;32(9):896–902. Epub 2014/08/24. doi: 10.1038/nbt.2931. PubMed PMID: 25150836.

69. Robinson MD, McCarthy DJ, Smyth GK. edgeR: a Bioconductor package for differential expression analysis of digital gene expression data. Bioinformatics (Oxford, England). 2010;26(1):139–40. Epub 2009/11/11. doi: 10.1093/bioinformatics/btp616. PubMed PMID: 19910308.

70. Chen J, Bardes EE, Aronow BJ, Jegga AG. ToppGene Suite for gene list enrichment analysis and candidate gene prioritization. Nucleic Acids Res. 2009;37(Web Server issue):W305-W11. Epub 2009/05/22. doi: 10.1093/nar/gkp427. PubMed PMID: 19465376.

71. Brewer GJ, Torricelli JR. Isolation and culture of adult neurons and neurospheres. Nature Protocols. 2007;2(6):1490–8. doi: 10.1038/nprot.2007.207.

72. Stuart T, Butler A, Hoffman P, Hafemeister C, Papalexi E, Mauck WM, 3rd, et al. Comprehensive Integration of Single-Cell Data. Cell. 2019;177(7):1888-902.e21. Epub 2019/06/11. doi: 10.1016/j.cell.2019.05.031. PubMed PMID: 31178118; PubMed Central PMCID: PMCPMC6687398.

73. Hum NR, Sebastian A, Gilmore SF, He W, Martin KA, Hinckley A, et al. Comparative Molecular Analysis of Cancer Behavior Cultured In Vitro, In Vivo, and Ex Vivo. Cancers (Basel). 2020;12(3). Epub 20200314. doi: 10.3390/cancers12030690. PubMed PMID: 32183351; PubMed Central PMCID: PMCPMC7140030.

